# Microbial strong organic ligand production is tightly coupled to iron in hydrothermal plumes

**DOI:** 10.1101/2023.01.05.522639

**Authors:** Colleen L. Hoffman, Patrick J. Monreal, Justine B. Albers, Alastair J.M. Lough, Alyson E. Santoro, Travis Mellett, Kristen N. Buck, Alessandro Tagliabue, Maeve C. Lohan, Joseph A. Resing, Randelle M. Bundy

## Abstract

Hydrothermal vents have emerged as an important source of iron to seawater, yet only a subset of iron is soluble and persists long enough to be available for surface biological uptake. The longevity and solubility of iron in seawater is governed by strong organic ligands, like siderophores, that are produced by marine microorganisms and are a part of the ocean’s dissolved iron-binding ligand pool. These ligands have been hypothesized to aid in the persistence of dissolved iron in hydrothermal environments. To explore this hypothesis, we measured iron, iron-binding ligands, and siderophores from 11 geochemically distinct sites along a 1,700 km section of the Mid-Atlantic Ridge. Siderophores were found in hydrothermal plumes at all sites, with proximity to the vent playing an important role in dictating siderophore types and diversity. The notable presence of amphiphilic siderophores may point to microbial utilization of siderophores to access particulate hydrothermal iron, and the exchange of dissolved and particulate iron. The tight coupling between strong ligands and dissolved iron within neutrally buoyant plumes across six distinct hydrothermal environments, and the presence of dissolved siderophores with siderophore-producing microbial genera, suggests that biological production of siderophores exerts a key control on hydrothermal dissolved iron concentrations.

## 1 Introduction

Over the last few decades, observations and modelling efforts have increased our understanding about the critical role organic ligands play in the cycling, transport, and utilization of trace metals (Tagliabue et al., 2017; Buck et al., 2018; Bundy et al., 2018; Moore et al., 2021). Strong iron-binding organic ligands (L_1_) are a heterogeneous mixture of microbially produced compounds that are operationally classified based on their binding strength with iron (Fe) 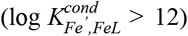, and are thermodynamically favored to complex and stabilize external sources of Fe to prevent its scavenging and removal. For example, in high dissolved and particulate Fe estuarine systems, only the dissolved Fe (dFe) bound to the strongest Fe-binding ligands remains in solution (Bundy et al., 2015; Buck et al., 2007) and is accessible for downstream biological uptake.

Siderophores are the strongest known Fe-binding organic ligands. They are produced by bacteria and fungi to facilitate Fe uptake and solubilization of otherwise inaccessible phases in the marine environment (Butler, 2005; Manck et al., 2022). They have primarily been considered as an important microbial strategy for Fe acquisition in the low-Fe surface ocean (Vraspir and Butler, 2009; Butler, 2005). However, siderophore uptake and biosynthesis genes were observed in >70% of Fe-related bacterial transcripts at Guaymas Basin (Li et al., 2014), have been identified in oxygen-deficient zones (Moore et al., 2021), and is a common Fe acquisition strategy within terrestrial and pathogenic ecosystems (Sandy and Butler, 2009) where Fe is orders of magnitude higher than seawater.

Although other unknown strong Fe-binding ligands have been observed in hydrothermal plumes (Buck et al., 2018) and throughout the deep ocean as well as siderophores observed below the euphotic zone (Bundy et al., 2018), no previous studies have ever directly characterized siderophores in hydrothermal systems. Some form of ‘stabilizing agent’ (i.e. ligands) has been proposed for the long-range transport of hydrothermal Fe into the ocean interior. The role of strong Fe-binding ligands in hydrothermal dFe transport represents an important gap knowledge gap with how hydrothermal vents may impact the ocean dFe inventory (Resing et al., 2015). Here, for the first time, we identified siderophores and siderophore-producing microbes in 11 geochemically distinct hydrothermal environments along the slow-spreading (20-50 mm/yr) Mid-Atlantic Ridge (MAR), including four black smokers (high temperature, high Fe), four off-axis sites, one diffuse vent (low temperature, low Fe), one alkaline vent (pH 9-11, very low Fe), and one non-vent fracture zone, to understand whether microbial ligand production impacts the supply of hydrothermal dFe to the ocean. Overall, our results show microbially-produced siderophores were present in all sites, and that strong L_1_ ligands are tightly coupled to hydrothermal dFe in this system. Strong organic ligands produced by bacteria thus play a key role in deep ocean Fe delivery from hydrothermal systems.

## 2 Results and Discussion

### 2.1 The role of iron-binding ligands and siderophores in hydrothermal plumes

Strong organic Fe-binding ligands, or L_1_ ligands, are important for stabilizing Fe in hydrothermal plumes (Tagliabue et al., 2017; Resing et al., 2015; Buck et al., 2018). The average binding strength and concentration of organic Fe-binding ligands were quantified in multiple vent systems that spanned a wide range in dFe concentrations (0.41-90.3 nM) and underlying vent geology. Over 99% of dFe in the neutrally buoyant plumes were complexed by L_1_ ligands and the ligands were almost always completely saturated with dFe, meaning Fe-free ‘excess’ L_1_ ligands capable of binding additional Fe were present in low concentrations (< 1 nM; **Fig. S1**). As a result, dFe concentrations were tightly coupled to L_1_ ligands in a nearly 1:1 ratio (**Fig. 1d**), similar to previous studies in other neutrally buoyant plumes (**Fig. 1e**) (Lough et al., 2022; Buck et al., 2018, 2015). The strong coupling between dFe and ligands was only observed at sites where L_1_ ligands were detected. Some sampling locations, such as in the buoyant plume, contained high concentrations of weaker ligands (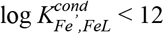, **Table S2**) with no correlation to dFe.

**Figure 1.**
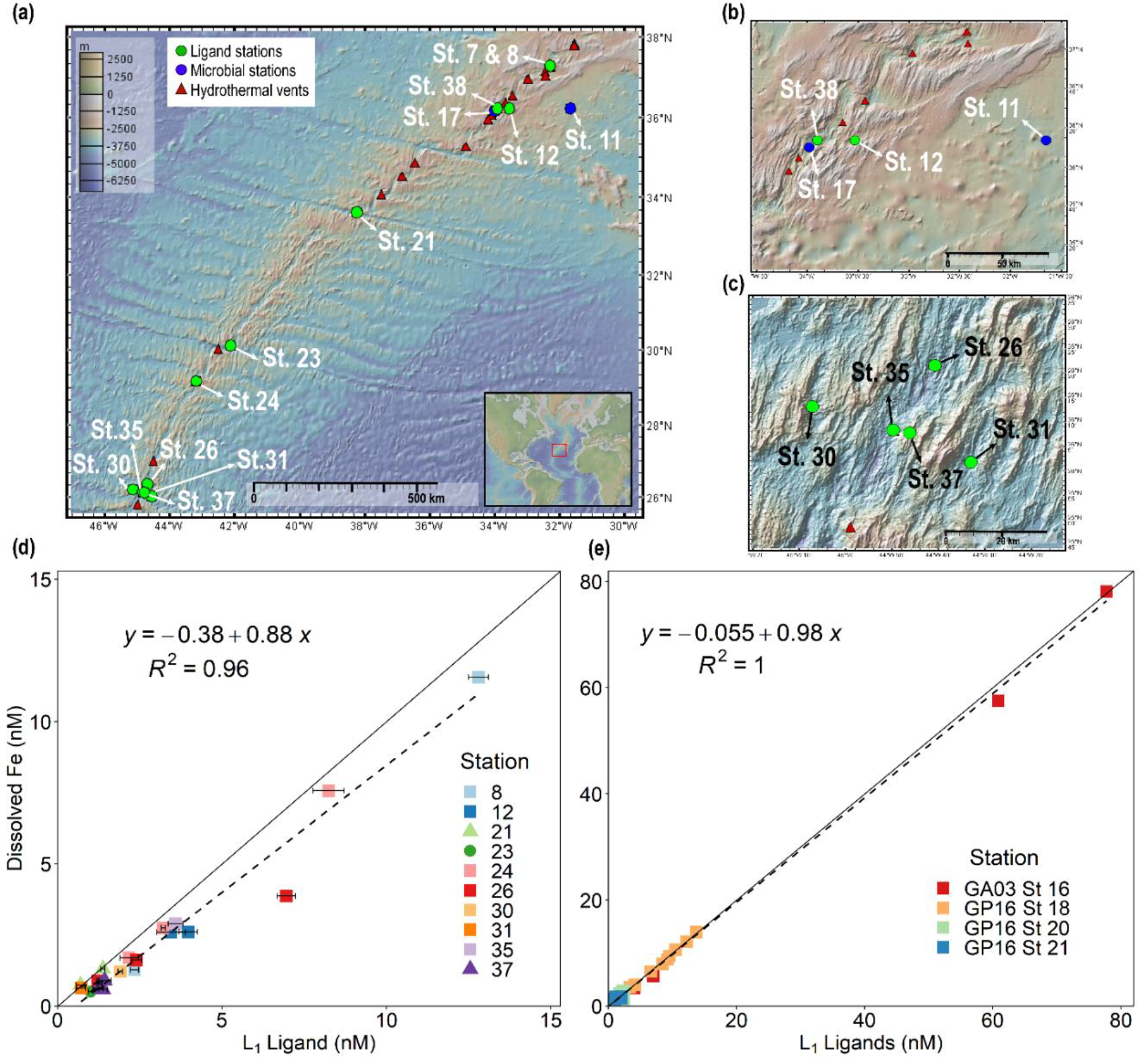
Dissolved iron is strongly correlated with L_1_ iron-binding ligands in diverse hydrothermal systems. (a) Station map showing the 11 sites investigated along the MAR. Known hydrothermal vents are marked as red triangles(Beaulieu and Szafrański, 2020). Two expanded inset maps for (b) Rainbow and (c) TAG hydrothermal vent fields. For additional information about vent site characteristics refer to **Table 1**. (d) dFe versus L_1_ iron-binding ligands at each vent site in this study showing a ∼1:1 correlation (m= 0.88, R^2^ = 0.96) with dFe in neutrally-buoyant plumes along the MAR. (e) dFe versus L_1_ ligands from previous studies over the ridge axis and ∼80 km from ridge axis in the Southern East Pacific Rise hydrothermal plume(Buck et al., 2018), and over TAG hydrothermal vent field(Buck et al., 2015). The solid black lines in (d) and (e) are the 1:1 ratio line between dFe and ligand concentrations, and dashed lines show the linear regression for the corresponding data. Square symbols refer to spreading centers, triangles refer to fracture zones, and circles refer to alkaline vents. Error bars represent the 95% confidence interval of the data fit as calculated by ProMCC(Omanović et al., 2015). The map was created using GeoMapApp version 3.6.14.

Our results indicate that L_1_ ligands uniquely set the dFe concentration in neutrally buoyant plumes. A similar control of dFe concentrations by L_1_ ligands has been previously observed in rivers (Buck et al., 2007) and aerosol solubility experiments (Fishwick et al., 2014). One explanation is that both the dFe and L_1_ ligands originate from the vent fluids themselves, yielding a tightly coupled hydrothermal endmember. However, the concentration of L_1_ ligands did not correlate with excess mantle Helium-3 (^3^He_xs_, **Fig S2, Table S2**) (Lough et al., 2022), a nearly conservative tracer of the mixing of hydrothermal fluids with seawater (Buck et al., 2018). These results suggest the L_1_ ligands were not sourced from the vent fluids along with dFe. All known sources of L_1_ ligands are biologically produced. Therefore, the L_1_ ligands observed here could be sourced either from bacteria that produced them in the surrounding deep ocean seawater that was then entrained, local production from vent-biota and/or microbial mats, diffusion from microbial production in sediments, or *in-situ* production by bacteria within the neutrally buoyant plume (Mellett et al., *submitted*).

Microbial organic ligand production is thought to tightly control global Fe inventories in both hydrothermal plumes (Cowen and Bruland, 1985; Cowen et al., 1990) and the open ocean (Lauderdale et al., 2020). Siderophores are operationally defined as L_1_ ligands by their binding strength (Moore et al., 2021; Manck et al., 2022) and have been proposed as important L_1_ ligands in hydrothermal plumes (Li et al., 2014), though they have never been directly measured. We used state-of-the-art liquid chromatography coupled to electrospray ionization mass spectroscopy (Boiteau et al., 2016) in a targeted approach to identify discrete components of the L_1_ ligands and to search for known siderophores. We observed a large diversity of siderophores with high confidence in every vent site using mass-to-charge ratio (*m/z*), MS/MS spectra, and specific chromatographic characteristics (**Fig. 2a**). Relative peak areas as a proxy for concentrations of putative siderophores also significantly correlated with dFe, as observed with dFe and L_1_ ligands (**Fig. 2b**).

**Figure 2.**
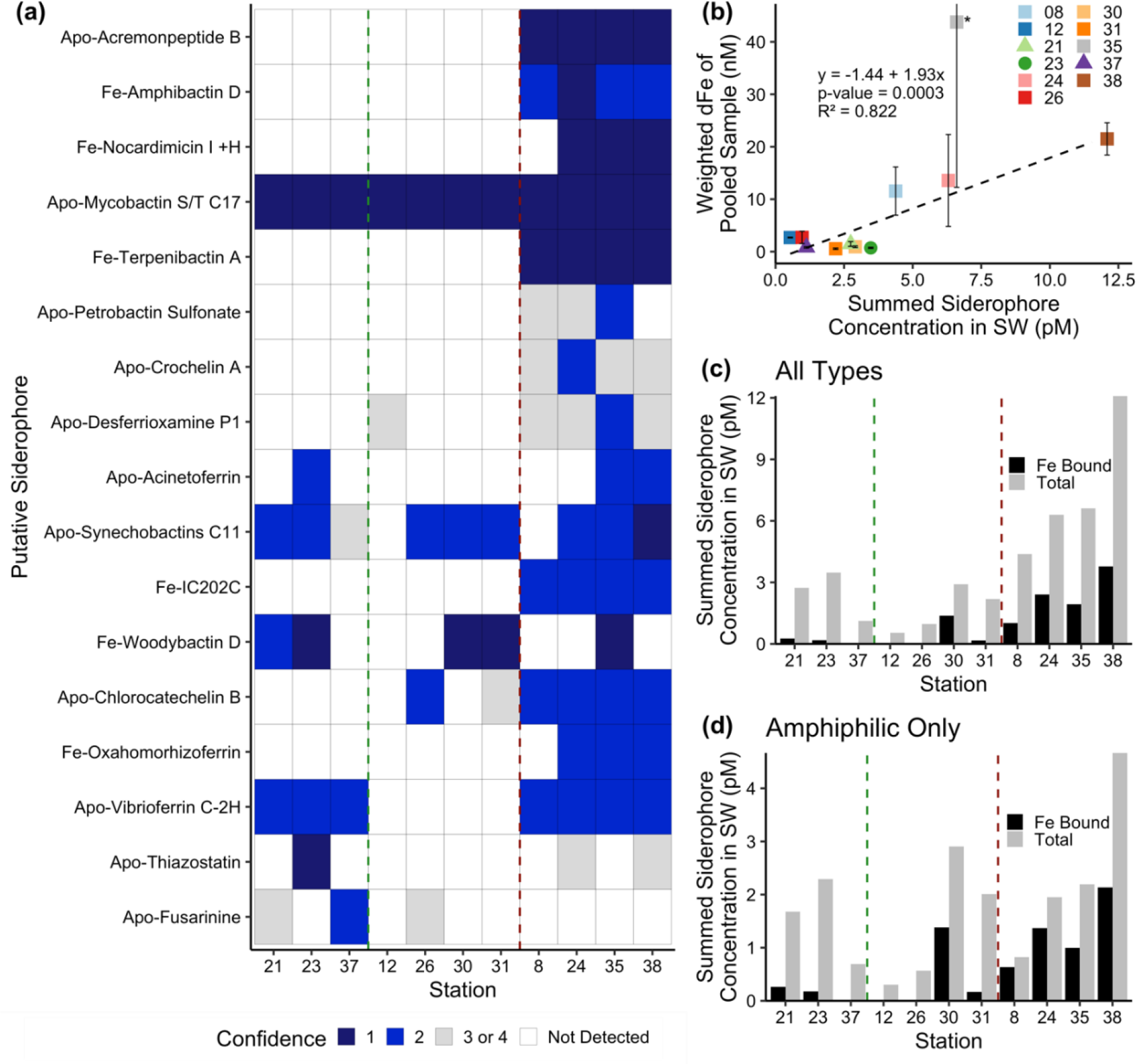
Siderophore presence in hydrothermal plumes along the MAR. (a) Heat map of confidence levels 1-2 (blue gradient, 1 = highest confidence). Gray boxes indicate a detection with lower confidence (see Methods), and white boxes indicate no detection at those sites. The y-axis is ordered from top to bottom in terms of descending mass of the apo (without Fe) form of the siderophore. (b) Model II ordinary least squares regression on dFe versus summed siderophore concentrations (of detections in Fig. 2b), calculated from peak areas, at each site. Since the siderophore analysis was performed on pooled samples, the dFe values in the regression are weighted values based on measured dFe and volume of each constituent of the pooled sample. The vertical error bars represent the standard deviation of dFe of the constituents. TAG (St. 35) was not included in the regression due to its large range of dFe values and outlier behavior. (c-d) Fe bound versus total summed concentration of (c) all types of siderophores and (d) amphiphilic siderophores at each station. The vertical green lines separate fracture/diffuse sites from off-axis sites and vertical red lines separate off-axis from on-axis sites as defined in Table 1. Symbols follow Fig. 1.

Siderophores were present in concentrations similar to the surface ocean (Boiteau et al., 2016; Moore et al., 2021; Park et al., 2022; Bundy et al., 2018), and comprised 0.01-0.4% of the total L_1_ ligands (**Table 1**). This is likely a significant underestimate of siderophore contributions to the L_1_ ligand pool due to analytical constraints in identifying unknown siderophores. That is, known siderophores represent a small fraction of what is expected to be produced in nature (Hider and Kong, 2010). In addition, most siderophores are not commercially available to use as standards, and individual siderophores have different ionization or extraction efficiencies. We also restricted our reporting to compounds only identified with very high confidence (**Fig 2a, S3**). The extraction efficiency for the solid phase extraction technique is around 10% for bulk Fe-binding organics (Bundy et al., 2018) and 40% for a siderophore standard (Waska et al., 2015). Correcting for extraction efficiency of the identified siderophores yields contributions of 1-4% of the L_1_ pool. We are inevitably missing other naturally occurring unknown compounds. Regardless of the small percentage contribution to total L_1_ ligands, it is evident that microbially produced siderophores are ubiquitous across all vent sites.

**Table 1.**
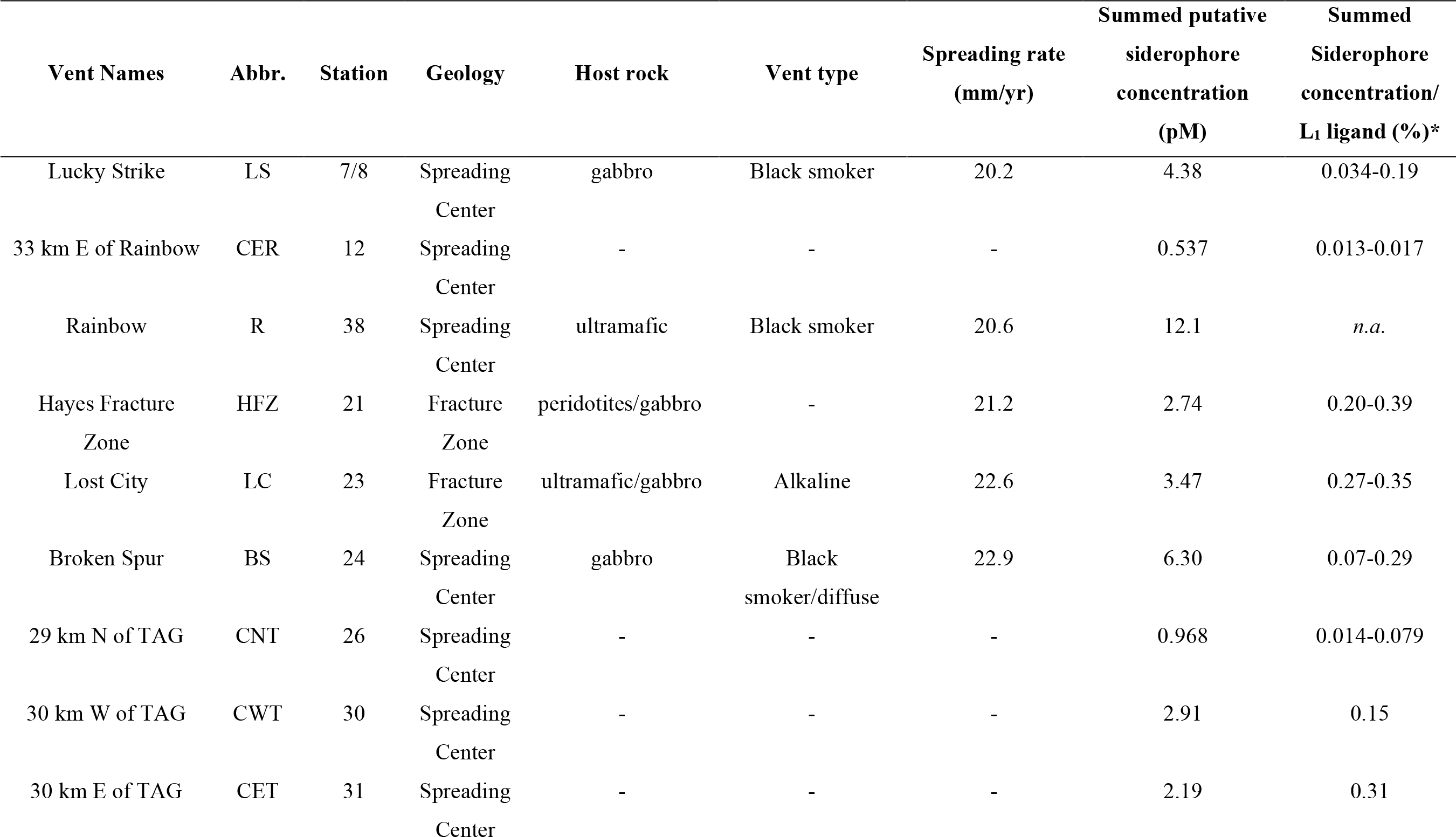
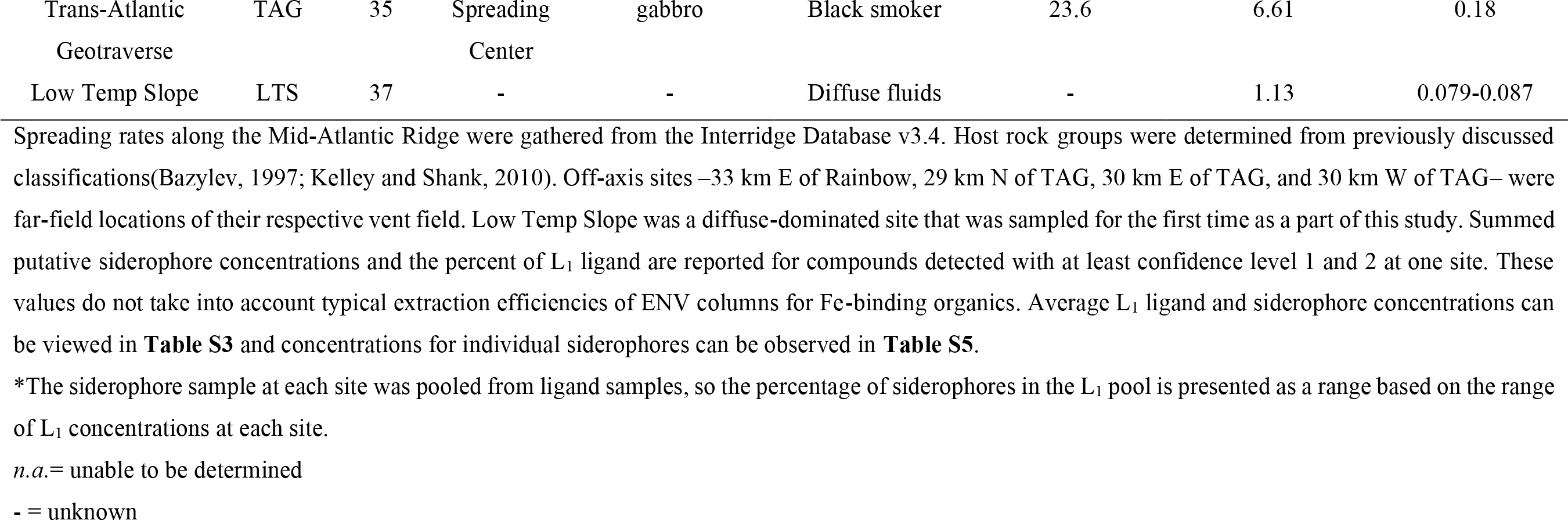
Characteristics of sample locations along the Mid Atlantic Ridge.

The high diversity of siderophores across a huge range of hydrothermal vent systems reveals several surprising aspects of Fe cycling. The biosynthesis of a siderophore is energy-intensive and is regulated by Fe concentration in the surrounding environment. Siderophore presence suggests that bacteria are producing these compounds despite the overall higher Fe concentrations in the deep ocean and in hydrothermal plumes. Consistent with siderophore utilization in terrestrial ecosystems, one hypothesis is that siderophore production is beneficial to bacteria in the plumes for transforming Fe from otherwise inaccessible forms, such as particulate nanopyrites or Fe oxyhydroxides. The identification of siderophores — and their relationship with dFe — provides compelling evidence that microbial production of ligands is responsible for some portion of the tight coupling between L_1_ and dFe in hydrothermal systems along the MAR.

### 2.2 Identity of siderophores in hydrothermal systems

Marine ligand composition changes with environmental gradients (Boiteau et al., 2016), making the structure and functional groups of siderophores identified in and around hydrothermal samples of particular interest. Samples collected from on-axis spreading centers contained the highest dFe concentrations (> 20 nM) and wider variety of siderophores than samples from fracture zones, diffuse, and off-axis sites (dFe ≤ 1 nM). The greatest number of distinct siderophores were identified at Lucky Strike, Broken Spur, Rainbow, and TAG (**Fig. 2**). On average, 13 compounds were identified with high confidence per on-axis spreading center sample, compared with 5 per diffuse/fracture zone sample, and 2.5 per off-axis sample (**Fig. 2b, Fig. S4**). Mixed-type siderophores — containing different moieties that bind to Fe(III) — were common at all sites. Hydroxamates were identified at and around spreading centers, yet none of these were detected with high confidence in samples from diffuse/fracture zones (**Fig. S4**). Vent type and proximity played a role in the diversity and abundance of siderophore types produced, likely related to the diversity of the microbial community and/or unique Fe acquisition strategies across sites.

Low Fe surface waters in the ocean have higher concentrations of amphiphilic siderophores compared to high Fe waters and terrestrial systems (Boiteau et al., 2016). Amphiphilic siderophores have long hydrocarbon tails that can be embedded into the lipid bilayer of the bacterial cell membrane providing a mechanism to shuttle Fe into the cell and prevent diffusive loss (Martinez et al., 2003). Amphiphilic siderophores accounted for 57% of the siderophores in our samples (**Fig. S5**), supporting the ubiquity of amphiphilic siderophores in the marine environment (Butler and Theisen, 2010). It was surprising they were so common, due to the elevated Fe concentrations observed relative to the Fe-poor surface ocean. Amphiphilic siderophores were found in concentrations between 0.3-4.7 pM, with highest found at Rainbow (**Fig. 2d, Table S5**). These concentrations were similar to those observed in the upper ocean (Boiteau et al., 2016; Bundy et al., 2018; Boiteau et al., 2019). Marine bacteria produce suites of amphiphilic siderophores as a way to adapt to the change in hydrophilicity in the surrounding environment (Sandy and Butler, 2009; Homann et al., 2009). Unlike in the surface ocean where amphiphilic siderophores are observed in Fe-limited regions (Boiteau et al., 2016), amphiphilic siderophores in plumes could be a way for bacteria to access Fe as they are physically transported and cope with strong chemical gradients, similar to the production of multiple siderophores in terrestrial and pathogenetic systems as a means to access inorganic particulate Fe for cellular uptake and storage (Hider and Kong, 2010).

### 2.3 Microbial sources of siderophores in hydrothermal plumes

Microbial production of siderophores is a strategy for organisms to adapt or compete with others for Fe (Sandy and Butler, 2009). We examined microbial community composition around Rainbow (St. 11, 17) and Lucky Strike (St. 7; **Table 1, Table S1**) using 16S rRNA gene-based amplicon sequencing to detect bacteria with the metabolic potential to synthesize siderophores (**Fig. 3, S11**), where the presence of taxa encoding siderophore biosynthetic gene clusters indicates whether the microbial community is genetically capable of producing the compounds we observed. Bacterial genera containing known siderophore-producers were found at all three MAR sites examined, and putative siderophore-producers represented 3-20% of the relative abundance of each community (**Fig. 3**). Putative siderophore-producers were more abundant in the 3 μm (particle-attached) size fraction than in the 0.2 μm (free-living) fraction, suggesting siderophore production is more common in particle-associated bacteria in hydrothermal environments.

**Figure 3.**
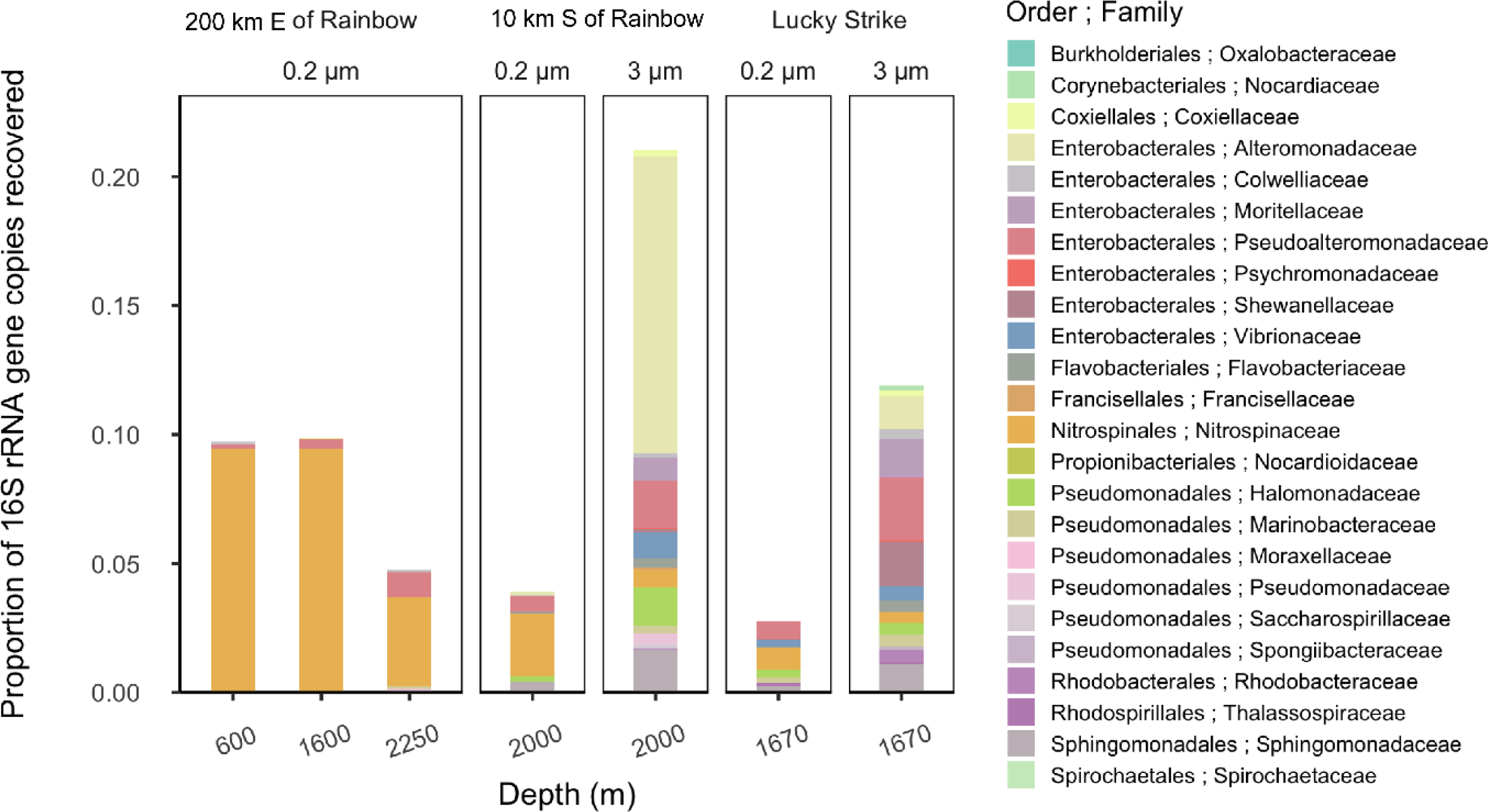
Relative abundance of putative siderophore-producing taxa. Bar height indicates the proportion of 16S rRNA genes recovered in each sample, separated by depth, filter size fraction, and site location. Colors correspond to taxonomy. Genera found in MAR vent microbial communities with members in the antismash database predicted to produce siderophores are depicted at the family level.

We found microbial genera in our samples that can produce a subset of the siderophores identified here, including ferrioxamines, vibrioferrin, and acinetoferrin (Butler, 2005; Vraspir and Butler, 2009; Moore et al., 2021; Bundy et al., 2018; Boiteau et al., 2016). Genera with the genetic potential to produce ferrioxamines were present at all three sites, while those known to produce vibrioferrin were present at Lucky Strike and Rainbow, and those producing acinetoferrin were also present at Rainbow (**Table S1, S6**). Mycobactins were detected with high confidence in every sample of this study, and genes encoding mycobactin have been detected in a cultured organism from a hydrothermal system (Gu et al., 2019), but no mycobactin producers were identified in this study. We detected woodybactin D with high confidence in 5 out of 11 sites. Although these biosynthetic genes were not identified in any of the genera observed, woodybactin D is a carboxylate siderophore isolated from *Shewanella* (Carmichael et al., 2019), and groups of deep-sea *Shewanella* (Kato and Nogi, 2001) were found in the dataset (**Fig. S11**). The biosynthesis genes for many of the siderophores identified are unknown. Thus, finding genera capable of producing only a subset of the siderophores characterized is not surprising. The observation that a significant portion of the *in-situ* microbial community is capable of synthesizing siderophores (**Fig 3**) suggests that siderophore production is more widespread in the marine environment and deep ocean than previously believed.

Although siderophores are often assumed to be associated primarily with low Fe conditions, evidence that siderophores are ubiquitous in the marine environment — including higher Fe ones — has been increasing (Park et al., 2023). The higher dFe associated with hydrothermal plumes may still not be high enough to suppress siderophore production due to the Fe requirements of bacteria (Tortell et al., 1996). It is also likely that in hydrothermal plumes not all of the Fe is bio-accessible. For example, soil microbes secrete siderophores to solubilize particulate Fe (Crowley et al., 1991). Similar processes could be occurring in hydrothermal plumes, where Fe mineral phases are common (Hoffman et al., 2020). It is telling that most siderophore-producing genera were found to be particle-associated (**Fig. 3**), providing additional evidence that siderophores might be produced to solubilize particulate Fe. Further work that assesses why bacteria are producing siderophores in neutrally buoyant plumes will be important for understanding the role of these complexes in facilitating the exchange of Fe between dissolved and particulate phases (Fitzsimmons et al., 2017), and how Fe-siderophore complexes may be stabilizing dFe in hydrothermal plumes.

Organic Fe-binding ligands have been implicated in playing a critical role in the preservation and transport of hydrothermal Fe into the ocean interior (Hoffman et al., 2018; Resing et al., 2015; Fitzsimmons et al., 2017; Toner et al., 2009; Bennett et al., 2011, 2008; Buck et al., 2018; Sander and Koschinsky, 2011). In this work, L_1_ ligands were tightly coupled to dFe in neutrally buoyant hydrothermal plumes along the MAR and found to be microbially produced. For the first time, specific siderophores were identified and demonstrated a bacterial source of L_1_ ligands being produced in response to external hydrothermal Fe inputs. Most of the bacteria putatively capable of synthesizing siderophores were particle-associated. This adds to a growing body of evidence that bacteria are using L_1_ ligands or siderophores to access particulate Fe pools, which is a key mechanism for controlling dFe in neutrally buoyant plumes. Exploring whether the Fe-organic ligand production is tightly coupled across additional hydrothermal vent systems will aid in constraining the biogeochemical importance of microbial feedbacks in impacting the hydrothermal dFe supply to the deep ocean.

## Supporting information

Supplemental Information_Hoffman_FRidge_revised

## Appendix Materials and Methods

### Sampling and cruise transect

Samples were collected as part of the 2017-2018 U.K. GEOTRACES GA13 section cruise along the Mid-Atlantic Ridge. Water samples from 11 venting and near venting locations were collected using a Seabird 911 conductivity, temperature, and depth (CTD) titanium rosette using conducting Kevlar wire with an oxidation-reduction potential (ORP) sensor to detect plumes. Teflon coated OTE (Ocean Test Equipment) bottles were pressurized to approximately 7 psi with 0.2 μm filtered air using an oil free compressor. A Sartobran 300 (Sartorius) filter capsule (0.2 μm) was used to collect filtered seawater samples into clean 250 mL LDPE sample bottles. Bottles and caps were rinsed 3 times with the filtered sample before being filled. Samples were stored frozen at -20°C for Fe-organic ligand characterization by voltammetry and mass spectrometry.

### Fe-binding ligand concentration and binding strengths Competitive Ligand Exchange-Adsorptive Cathodic Stripping Voltammetry

Fe-binding ligand concentrations and binding strengths were determined by competitive ligand exchange-adsorptive cathodic stripping voltammetry (CLE-ACSV) with a BAS*i* controlled growth mercury electrode (CGME) with an Ag/AgCl^-^ reference electrode and platinum auxiliary electrode (Bioanalytical Systems Incorporated). Using previously established methods (Buck et al., 2015, 2018; Bundy et al., 2018; Abualhaija and van den Berg, 2014; Hawkes et al., 2013b), 40 frozen filtrate (<0.2 μm) samples with dFe concentrations between 0.41-11.67 nM (**Table S1-S2**) were thawed in a 4°C fridge prior to analysis. A 15-point titration curve was analyzed for each sample. Briefly, within each titration, every point sequentially received 10 mL of sample, 7.5 μM of borate-ammonium buffer, 10 μM salicylaldoxime (SA) added ligand, and a dFe addition. Data was collected using the *Epsilon Eclipse Electrochemical Analyzer* (v.213) with a deposition time of 120 seconds and analyzed using *ElectroChemical Data Software* (v2001-2014) and *ProMCC* (v2008-2018) to determine peak areas and Fe-binding ligand parameters, respectively.

### Reverse Titration-CLE-ACSV

Reverse titration-CLE-ACSV (RT-CLE-ACSV) (Hawkes et al., 2013a) was completed on 10 samples from Broken Spur, and TAG hydrothermal vent fields with dFe concentrations between 19.01-90.25 nM (**Table S1-S2**). Briefly, a 10-point titration curve was analyzed for each sample with each titration point consisting of 10 mL of sample buffered with 7.5 μM boric acid and the competitive ligand 1-nitroso-2-napthol (NN) additions. All samples were analyzed on a BAS*i* Controlled Growth Mercury Electrode (CGME) with the *Epsilon Eclipse Electrochemical Analyzer* (v.213) and deposition time of 120 seconds. For each sample, competitive ligand NN additions were 0.5, 1, 2, 3, 4, 6, 9, 15, 20, and 40 μM. Samples were equilibrated overnight and purged with N_2_ (99.99%) for 5 minutes before analysis. At the end of each titration, three Fe additions (3-15 nM) were added to the final titration point to get the total concentration of Fe in equilibrium with ligands. Data was analyzed using *ElectroChemical Data Software* (v2001-2014) to acquire peak areas and a package in R using the model parameters of β_FeNN3_ = 5.12 x 10^16^, χ_min_ = 0.8, χ_max_ = 0.9, and *c1high* = 0.75 to determine the Fe-binding ligand parameters (Hawkes et al., 2013a). These parameters were chosen based on the recommendations for undersaturated samples and titrations curves where *ip*_*max*_ was not reached (Hawkes et al., 2013a). All other parameters within the model we kept at the default values.

### Fe-binding organic ligand quantification and characterization

In addition to determining the total concentrations of strong Fe-binding ligands (L_1_), we also identified and quantified siderophores that contributed to the L_1_ ligand pool. Between 0.65-1.5 L of 0.2 μm filtered seawater pooled from ligand samples at each site (described above) was pumped slowly (15-20 mL min^-1^) onto a polystyrene-divinylbenzene (Bond Elut ENV) solid phase extraction (SPE) column (Bundy et al., 2018; Boiteau et al., 2016). SPE columns were rinsed with MilliQ and stored at -20ºC until analysis. For the analytical measurements, samples were thawed in the dark, eluted in 12 mL of distilled methanol, and dried down to between 0.2-0.5 mL of sample eluent (**Table S1**). Aliquots were analyzed by reverse-phase liquid chromatography (LC) on a trace metal clean bio-inert LC (Thermo Dionex 3000 NCS). The LC was interfaced with an electrospray ionization-mass spectrometer (ESI-MS; Thermo Q-Exactive HF) to identify and quantify the compounds based on accurate mass (MS^1^) and the fragmentation (MS^2^) data (Bundy et al., 2018; Boiteau et al., 2016). MSconvert (Proteowizard) was used to convert MS data to an open source mzxML format, and two stages of data processing were conducted using modified versions of previously reported R scripts (Bundy et al., 2018; Boiteau et al., 2016). In the first stage, mzxML files were read into R using new package “RaMS” (Kulmer and Ingalls, 2022), and extracted ion chromatograms (EICs) were generated for each targeted *m*/*z* of interest from an in-house database of siderophores. The *m*/*z* targets were the ionized apo, ^54^Fe-bound, and ^56^Fe-bound version of each siderophore, with a tolerance of 7.5 ppm.

Putative siderophore candidates were filtered through a series of hard thresholds, such that MS^1^ spectra were quality controlled to contain a minimum of 25 datapoints and the maximum intensity of each EIC was greater than 1e4 counts. Spectra meeting these criteria and containing either ^54^Fe-bound and ^56^Fe-bound *m/z* peaks within 30 seconds of each other or an apo peak were displayed for the user to further inspect peak quality and make the final decision of whether to move on to stage two of processing with a given siderophore candidate.

Stage two of processing extracted MS^2^ spectra of the apo and Fe-bound forms of candidate siderophores to compare with the predicted MS^2^ generated by *in silico* fragmenter MetFrag (Ruttkies et al., 2016). The *in silico* fragmenter feature was run with a tolerance of 10 ppm on “[M+H]^+^” and “[M+Na]^+^” modes. A confidence level of 1-4, from highest to lowest confidence, was then assigned to putative siderophores based on the following criteria: (1) peaks were present in MS^1^ and MS^2^ spectra, and at least one of the three most-intense MS^2^ fragments matched *in silico* fragmentation, (2) peaks were present in MS^1^ and MS^2^ spectra, and smaller-intensity fragments matched *in silico* fragmentation, (3) peaks were present in MS^1^ and MS^2^ spectra, but little to no fragments matched *in silico* fragmentation, and (4) nicely shaped peaks were identified in MS^1^ spectra but no MS^2^ spectra was collected (outlined in **Table S4**; example spectra in **Fig. S6-S9)**. The confidence levels were modelled after reporting standards for metabolite identification (Sumner et al., 2007). MetFrag pulls chemical structures from publicly-available databases like PubChem or COCONUT(Sorokina et al., 2021), which contain most, but not all variations of siderophores. As such, Fe-bound candidates were usually run against the apo form available in the database, and for siderophores with similar structures but variations in fatty chain length or double bond placement, sometimes only one parent structure was available.

A 5-point standard curve with known concentrations of siderophore ferrioxamine E was used for quantification of putative siderophores, with a limit of detection of 0.257 nM in the eluent (**Fig. S10**), or 0.07-0.21 pM in the sample depending on sample-to-eluent volume ratio at each site (**Table S1**). MS^1^ peaks were integrated for all putatively identified siderophores and peak areas were converted to concentration using the standard curve and the concentration factor of sample volume to eluent volume (**Fig. S10**). Commercial standards are not available for most siderophores, and different compounds have distinct ionization efficiencies in ESI-MS. Thus, the siderophore concentrations reported here are estimates of siderophore concentrations in these environments based on ferrioxamine E. Additionally, 1 mM of cyanocobalamin was added as an internal standard to each sample aliquot to address any changes in sensitivity during LC-ESI-MS runs. All putative siderophores that were identified with peak areas less than the detection limit were discarded, and all remaining putative compounds with at least confidence levels 1 and 2 at one site were included in the manuscript and are referred to as siderophores throughout. Siderophore identifications remain putative due to inherent uncertainty with assignments by mass, but the confidence levels were designed such that high confidence candidates contain siderophore-like moieties in their fragments. Limited sample volumes prevented analysis via LC-ICP-MS like previous studies, which, in addition to greater availability of commercial standard, would allow definitive identification in future work. Confidence level 3 and 4 putative siderophores are only included in the Supplementary Information (**Table S5**). In a final step of quality control, EICs for ^13^C isotopologues of candidates were inspected to verify matching peak structure.

### Microbial community analysis

Microbial community composition was assessed in neutrally buoyant plumes and near venting sites at three sites: Lucky Strike (Station 7; 1670 m), 10 km S of Rainbow (Station 17; 2000 m), and 200 km E of Rainbow (Station 11; 600 m, 1600 m and 2250 m). A range of 1-2 L of seawater were filtered by pressure filtration through sequential 25 mm membrane filters housed in polypropylene filter holders (Whatman SwinLok, GE Healthcare, Pittsburgh, Pennsylvania) using a peristaltic pump and silicone tubing. Samples first passed through a 3 μm pore-size polyester membrane filter (Sterlitech, Auburn, Washington) then onto a 0.2 μm pore-size polyethersulfone membrane filter (Supor-200, Pall Corporation, Port Washington, New York). Pump tubing was acid washed with 10% hydrochloric acid and flushed with ultrapure water between each sample. The filters were flash frozen in liquid nitrogen in 2 mL gasketed bead beating tubes (Fisher Scientific) at sea.

Nucleic acids (DNA) were extracted as described previously(Santoro et al., 2010), with slight modifications. Briefly, cells on the filters were lysed directly in the bead beating tubes with sucrose-ethylene diamine tetraacetic acid (EDTA) lysis buffer (0.75 M sucrose, 20 mM EDTA, 400 mM NaCl, 50 mM Tris) and 1% sodium dodecyl sulfate. Tubes were then agitated in a bead beating machine (Biospec Products) for 1 min, and subsequently heated for 2 min. at 99°C in a heat block. Proteinase K (New England Biolabs) was added to a final concentration of 0.5 mg/mL. Filters were incubated at 55°C for approximately 4 h and the resulting lysates were purified with the DNeasy kit (Qiagen) using a slightly modified protocol (Santoro et al., 2010). The purified nucleic acids were eluted in 200 μL of DNase, RNase-free water, and quantified using a fluorometer (Qubit and Quanti-T HS reagent, Invitrogen Molecular Probes).

The 16S rRNA gene was amplified in all samples using V4 primers (Apprill et al., 2015; Parada et al., 2016) (515F-Y and 806RB) following a previously established protocol (Stephens et al., 2020). Amplicons were sequenced using a paired-end 250bp run on an Illumina MiSeq 500 and demultiplexed by the UC Davis Genome Center. The resulting 16S rRNA amplicon sequences were filtered and trimmed using the DADA2 pipeline in R(Callahan et al., 2016). Taxonomic assignments were made with version 138.1 of the SILVA SSU database (Quast et al., 2013) (silva_nr99_v138.1_wSpecies_train_set.fa.gz ; doi:10.5281/zenodo.4587955; accessed March 2022). Chloroplast and mitochondrial sequences were filtered out of the dataset using the ‘phyloseq’ R package (v 1.38.0), after which samples had read depths ranging from 9375 – 65486 reads (average 28425 ± 20014 reads) and represented 1010 unique amplicon sequence variants (ASVs). Read counts were transformed from absolute to relative abundance and taxa were aggregated to the Family level. The ten most abundant families present in each sample were visualized using the ‘ggplot2’ package (v. 3.3.5).

In order to assess the potential of the observed prokaryotic taxa to produce siderophores, we downloaded all siderophore biosynthetic gene clusters (BGCs) in the antismash secondary metabolite database (*n* = 7909) and used text-string matching to compare genera containing these BGCs to the genera found in our 16S rRNA gene dataset(Blin et al., 2021). We cross-referenced the nomenclature of antismash-predicted siderophores with that of the siderophores identified by LC-ESI-MS in this study, accounting for minor differences in naming convention between the two databases, to determine if microbial community members present at each site were predicted to make any of the siderophores that were measured at that site. Station 38 and Station 12 were the closest sites with siderophore measurements for comparison against the taxonomic samples taken at 200 km E of Rainbow and 10 km S of Rainbow, respectively. Samples for microbial taxonomy and siderophore identity were taken from the same location at Lucky Strike and thus directly compared.

## Data Availability

The CSV data reported in this study has been deposited at Zenodo under the DOI: http://doi.org/10.5281/zenodo.7325154. The LC-ES-MS data has been deposited on Massive under the DOI: http://doi.org/doi.10.25345/C5V97ZW7N. Microbial 16S rRNA data have been deposited on GenBank under the accession number BioProject #PRJNA865382. All data is freely available on each of these data repositories.

## Acknowledgments

We acknowledge the captain and crew of the R/V *James Cook*, Chief Scientist Alessandro Tagliabue, and Noah Gluschankoff for supporting this work. This study was a part of the FeRidge project (GEOTRACES section GA13) which was supported by the Natural Environment Research Council funding (NERC United Kingdom Grants NE/N010396/1 to MCL and NE/N009525/1 to AT). The International GEOTRACES Programme is possible in part thanks to the support from the U.S. National Science Foundation (Grant OCE-1840868) to the Scientific Committee on Oceanic Research (SCOR). CLH was funded by JISAO/CICOES postdoctoral fellowship. PJM was funded through the NOAA Hollings Scholar summer program. JR was funded by NOAA Ocean Exploration and Research, NOAA Earth-Ocean Interactions programs at NOAA-Pacific Marine Environmental Labs, and JISAO/CICOES. Part of this work was carried out in the University of Washington TraceLab, which receives support from the M.J. Murdock Charitable Trust in conjunction with the University of Washington College of Environment, and the Pacific Marine Environmental Labs at the National Oceanic and Atmospheric Administration. Parts of this work was also carried out in Dr. Anitra Ingalls laboratory with the help of Laura Truxal and Dr. Jiwoon Park at the University of Washington-School of Oceanography.

## Author Contributions

Manuscript preparation, sample/data processing, CSV analysis, and interpretation (C.L.H.), manuscript preparation, LC-ESI-MS data analysis and interpretation (P.J.M.), microbial analysis and interpretation (J.B.A. and A.E.S.), dissolved iron and derived excess ^3^He_xs_ measurements, sample collection (A.J.M. L. and M.C.L.), microbial data collection and ligand data interpretation (T.M. and K.N.B.), and project design and planning, data interpretation, and mentoring (A.T., M.C.L., J.A.R., and R.M.B.). All authors were involved in editing manuscript.

## Competing Interest Statement

The authors declare no competing interests.

## Notes

http://doi.org/10.5281/zenodo.7325154

http://doi.org/doi.10.25345/C5V97ZW7N

### Competing Interest Statement

The authors have declared no competing interest.

### Summary of Updates

This version has been shorted to match the submission requirements of Biogeosciences Letters.

