## Supplemental Information_Hoffman_FRidge_revised for "Microbial strong organic ligand production is tightly coupled to iron in hydrothermal plumes"

### S1. Supporting Information Text

#### S1.1 Additional Methods

**Competitive Ligand Exchange-Adsorptive Cathodic Stripping Voltammetry.** Two sets of Fe additions were used when conducting forward titrations to ensure full titration of samples. Dissolved Fe in the 40 samples process via forward titration ranged from 0.4-11.67 nM. For Lost City samples and one sample from Close E of TAG with a dissolved Fe concentration of 0.41 nM, dissolved Fe additions were 0, 0, 0.1, 0.25, 0.5, 1, 1.25, 1.5, 2, 2.5, 3, 4, 5, 7.5, and 10 nM. For the other 33 samples where a forward titration was conducted (**Table S1**), dissolved Fe additions were 0, 0, 0.25, 0.5, 1, 1.25, 1.5, 2, 2.5, 3, 4, 5, 7.5, 10, and 15 nM. Each sample was then equilibrated for at least 12 hours before analysis on the *BASi*.

**S2. Supporting Information Figures**

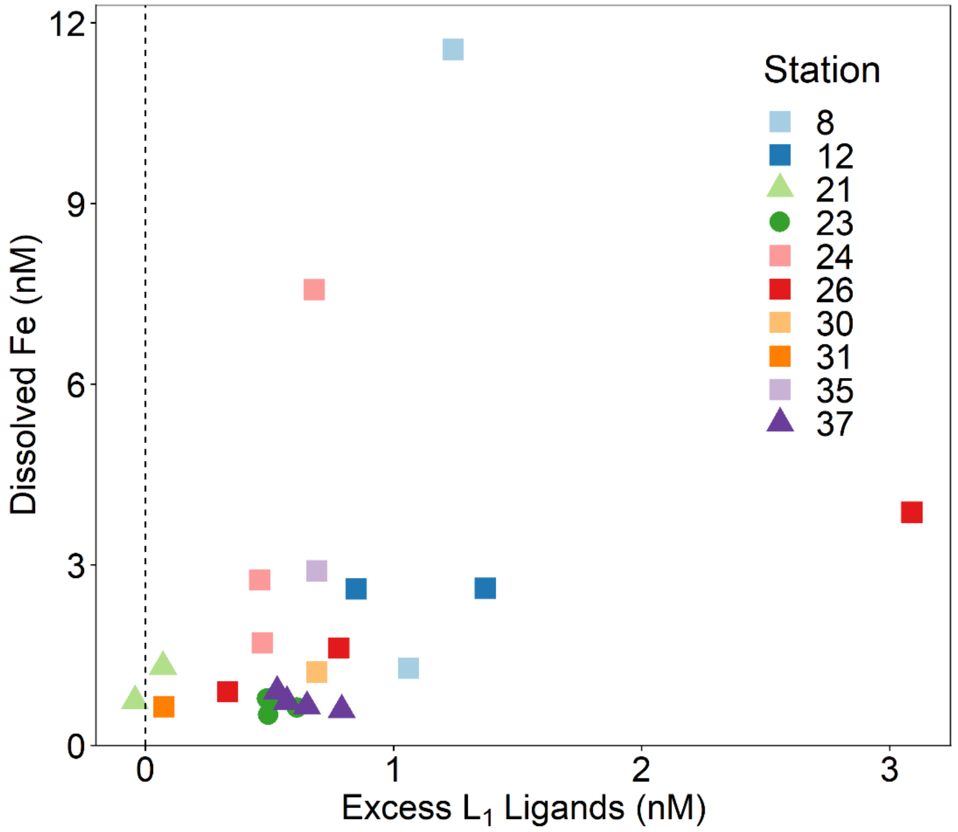

**Fig. S1. Excess $L_1$ along the Mid-Atlantic Ridge.** Excess $L_1$ ligands along the MAR. Hayes Fracture Zone (Station 21) was the only vent field to have one sample to not contain excess $L_1$ ligands. The dashed line is the zero line on the y-axis. Points with positive values correspond to samples with excess ligand present. Points with negative values correspond to samples with no excess ligand present. No $L_1$ or excess $L_1$ ligands were present at St. 38 Rainbow. Square symbols refer to spreading centers, triangles refer to fracture zones, and circles refer to alkaline vents.

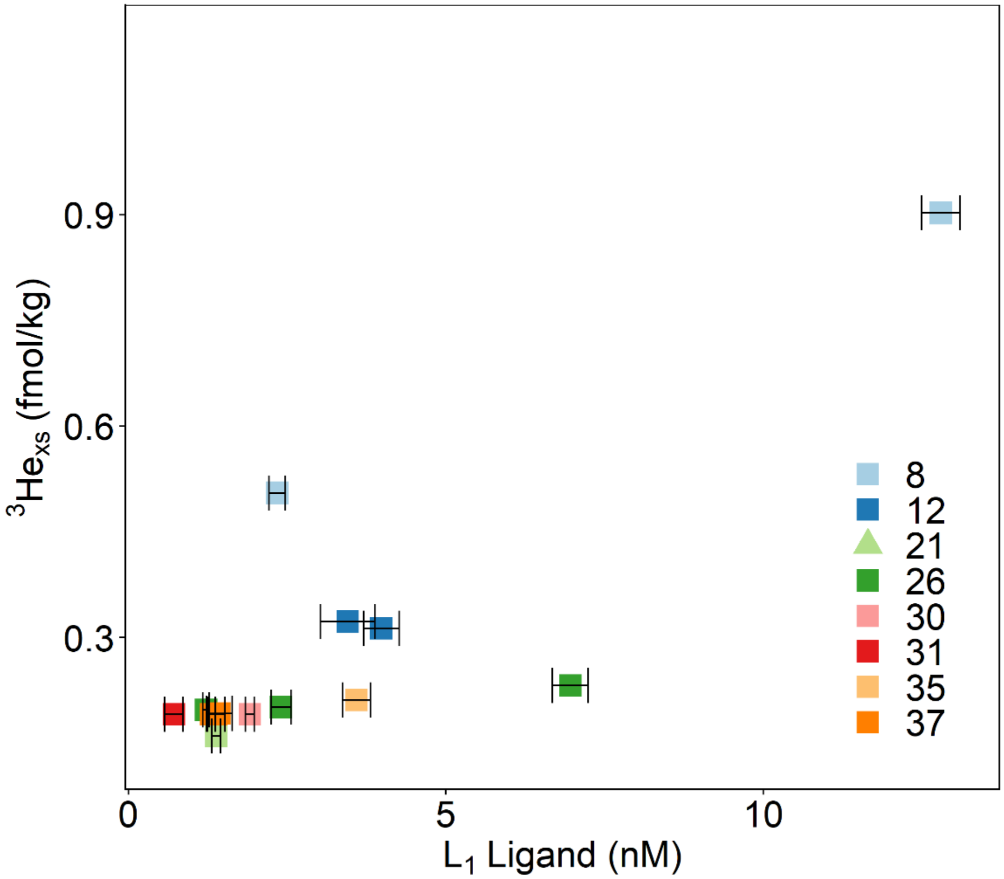

**Fig. S2. $L_1$ ligands versus $^3He_{xs}$ along the Mid-Atlantic Ridge.** $L_1$ ligands did not correlate with the $^3He_{xs}$ across the different plumes along the Mid-Atlantic Ridge. Expected $^3He_{xs}$ values were derived from the conservative relationship between dMn/$^3He_{xs}$ in hydrothermal plumes(1). No $^3He_{xs}$ was derived for St. 23 Lost City, St. 24 Broken Spur and one sample at St. 21 Hayes Fracture Zone (**Table S2**). Square symbols refer to spreading centers and triangles refer to fracture zones.

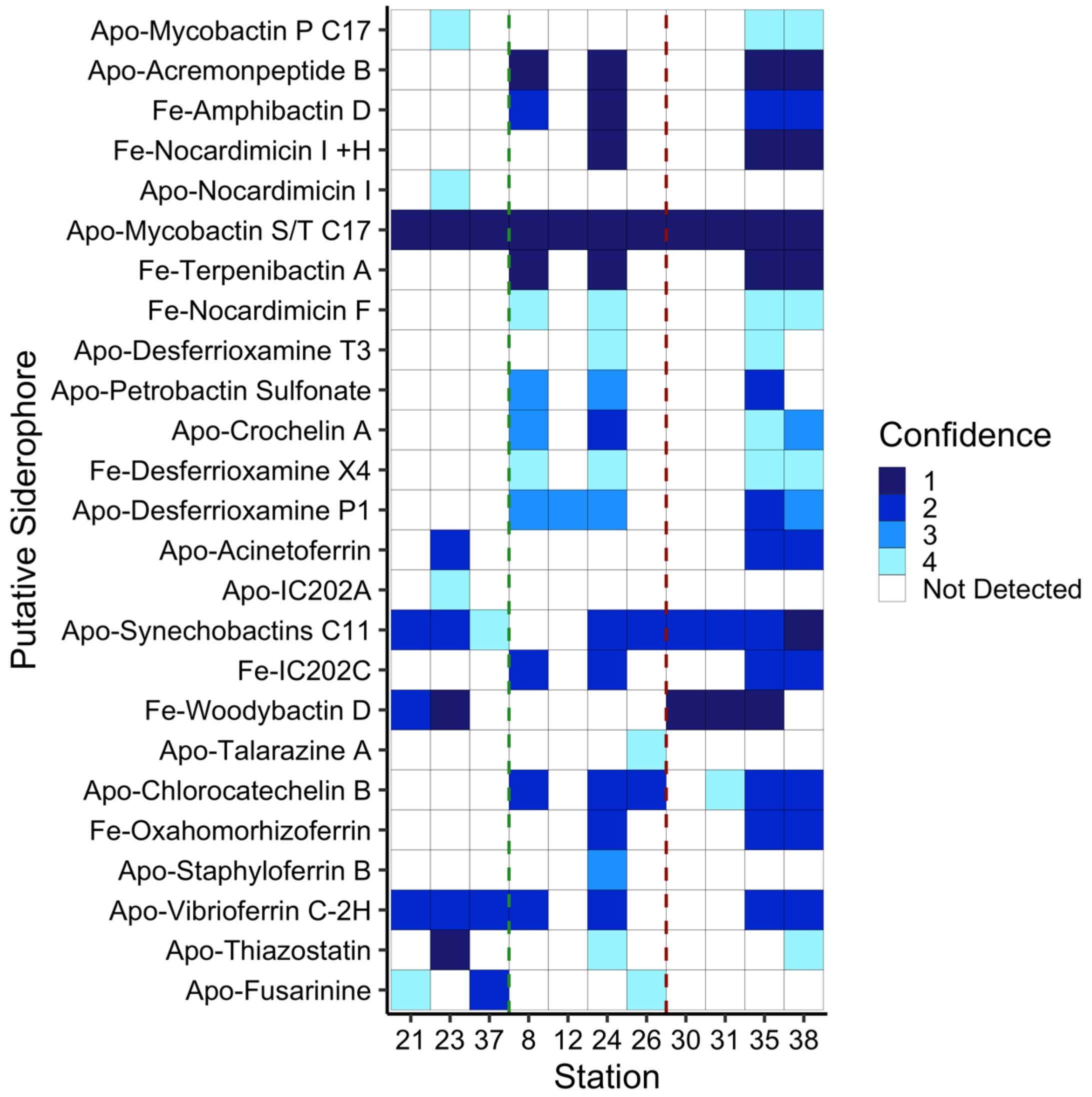

**Fig. S3. All putative siderophore identifications made in hydrothermal plumes along the MAR.** Heat map of siderophore detections at each site, similar to **Fig. 2a** of main text, but all siderophores and all confidence levels are visualized. The blue gradients indicates the confidence level of the identification, with level 1 representing the highest confidence level. In the text, confidence levels 1 and 2 are considered high confidence, while data from those only identified with confidence levels 3 and 4 not used in figures or statistics. Whites boxes indicate no detection at that particular site. The y-axis is ordered from top to bottom in terms of descending mass of the apo (without Fe) form. The vertical green lines separate fracture/diffuse dominant sites from off-axis sites, while the vertical red lines separate off-axis from on-axis sites. More information about the putative compounds are listed in **Table S5** and definitions of the confidence levels are found in the methods of the main text and are outlined in **Table S4**.

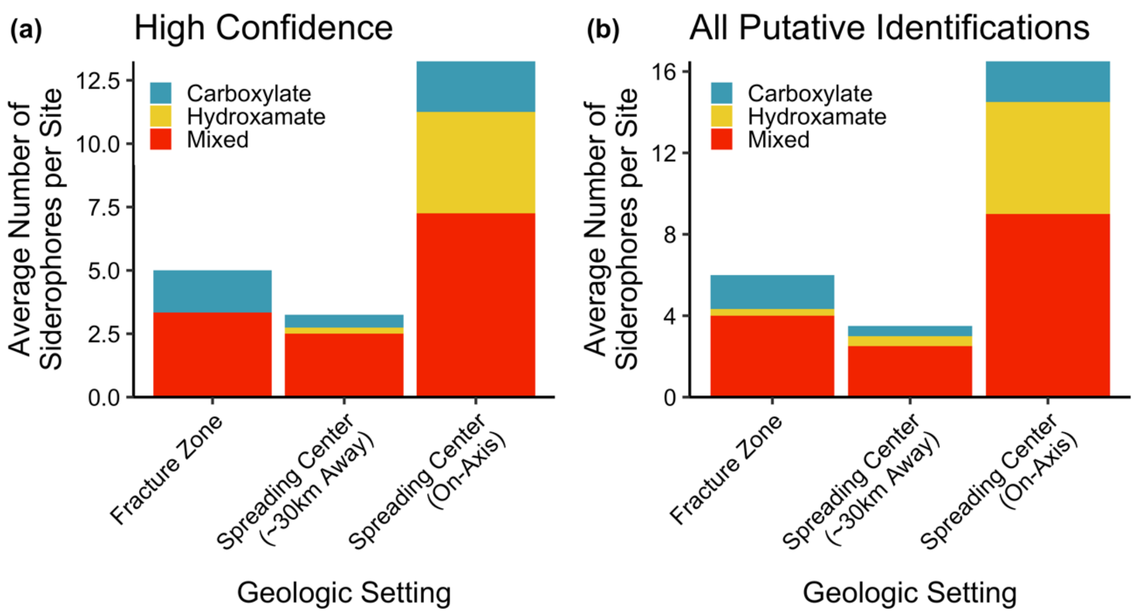

**Fig. S4. Distribution of types of siderophores.** Bar graph of types of putative siderophores identified with (a) high confidence (i.e. at least confidence level 1 and 2 at a site) and (b) all confidence levels, separated by the type of site. Sites above spreading centers averaged more putative siderophores than both those above fracture zone or lower temperature sites and those ~30km from the spreading center. Mixed-type siderophores dominated identification, with hydroxamate siderophores also significant above spreading centers.

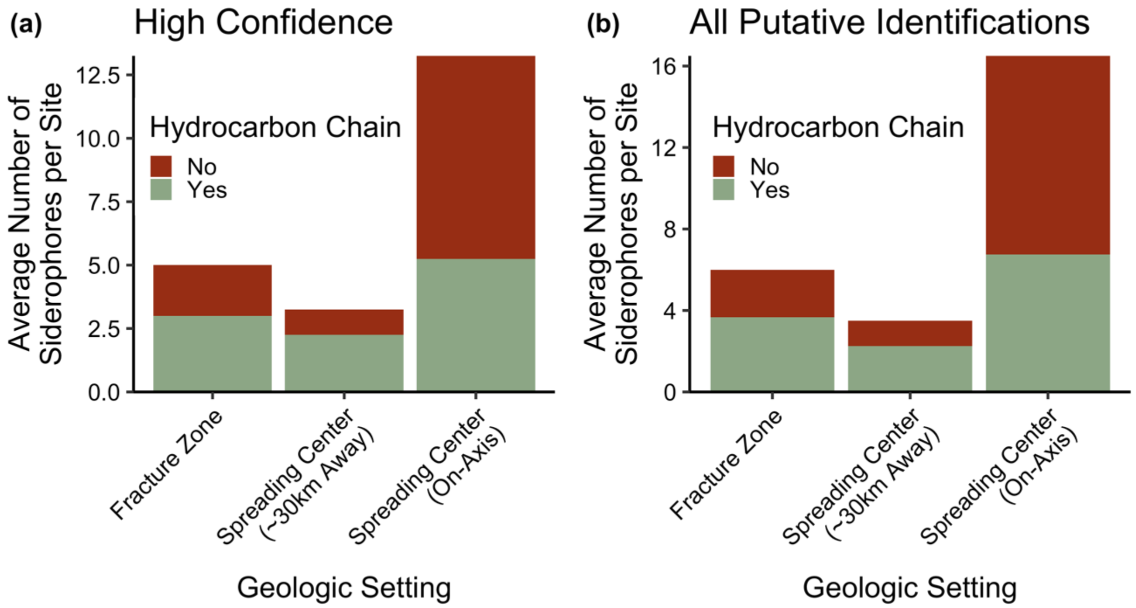

**Fig. S5. Distribution of amphiphilic siderophores.** Bar graph of types of putative siderophores identified with (a) high confidence (i.e. at least confidence level 1 and 2 at a site) and (b) all confidence levels, separated by the type of site. A "Yes" indicates that the structure of the siderophore contains a terminal hydrocarbon chain of 7+ carbon atoms. In total, over half (57%) of putative siderophores identified in this study contained a hydrocarbon chain.

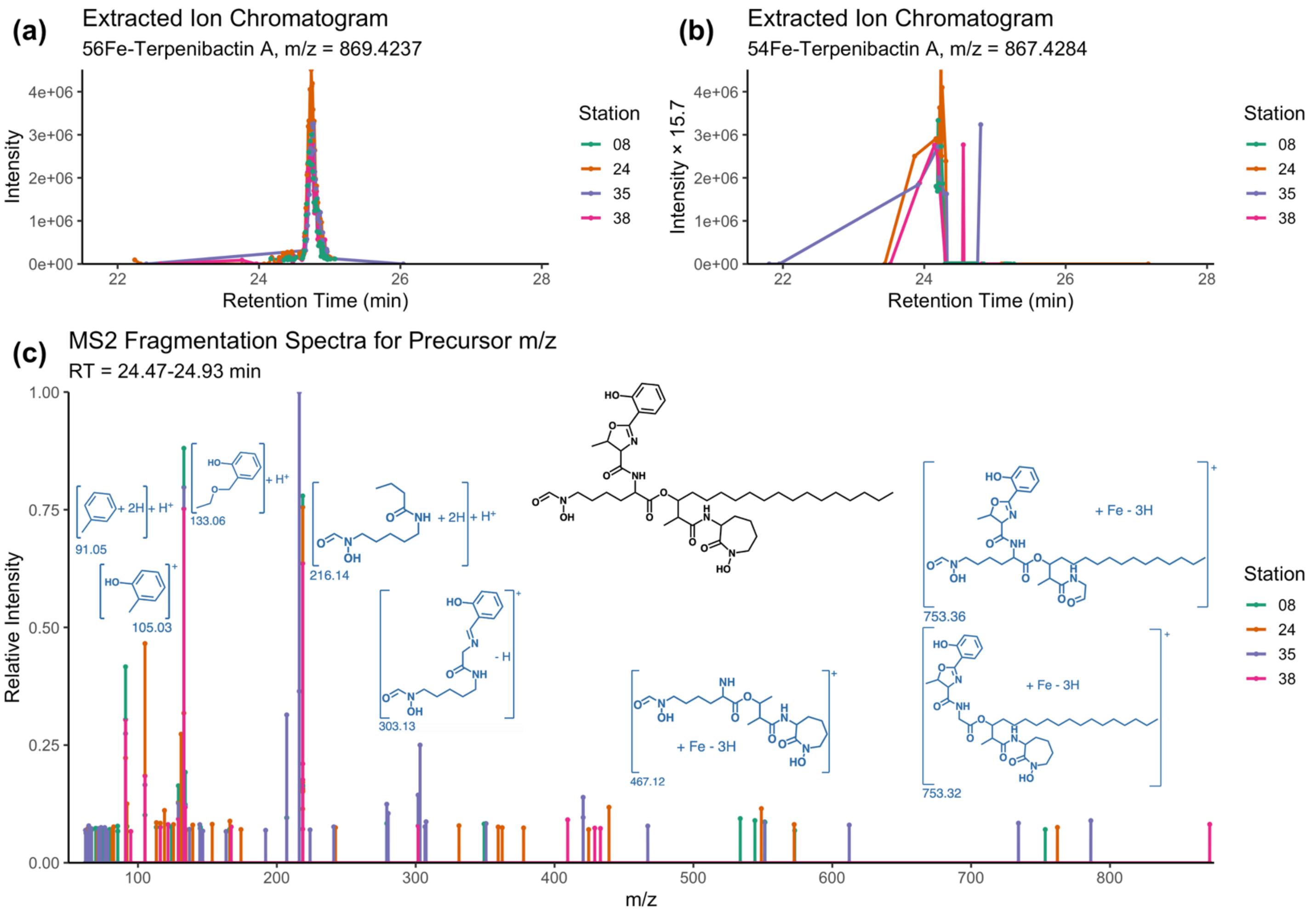

**Fig. S6. Confidence Level 1 example MS$^1$ and MS$^2$ spectra.** Example MS$^1$ (a-b) and MS$^2$ (c) spectra of putative siderophore Fe-Terpenibactin A is presented as an example of putative siderophores identified at the highest confidence level. The peak in the extracted ion chromatogram (EIC) corresponding the $^{56}$Fe-bound mass is presented in (a), while the EIC corresponding to the less-abundant $^{54}$Fe-bound mass is presented in (b), scaled by a factor of 15.7 (crustal abundance of $^{56}$Fe/$^{54}$Fe). Little is known about the isotopic fractionation associated with Fe-siderophore binding, but the fact that the $^{54}$Fe-Terpenibactin A signal scales to a similar intensity to that of $^{56}$Fe-Terpenibactin A provides strong evidence that the candidate compound is indeed bound to Fe. This strategy has been used in past siderophore identification workflows (2). Following analysis of the EICs, MS$^2$ fragments were extracted from the same *m/z* and retention time window and compared against *in silico* spectra for the candidate structure generated by MetFrag. For this particular compound, too many matching fragments were identified to display, but select fragments are presented in (c). The well-defined MS$^1$ peaks, consistent $^{54}$Fe and $^{56}$Fe EICs, and high-intensity fragments matching *in silico* fragmentation yields a detection with the highest confidence level. While this figure has spectra from multiple sites, confidence levels were typically assigned at each site independent of fragments at other sites.

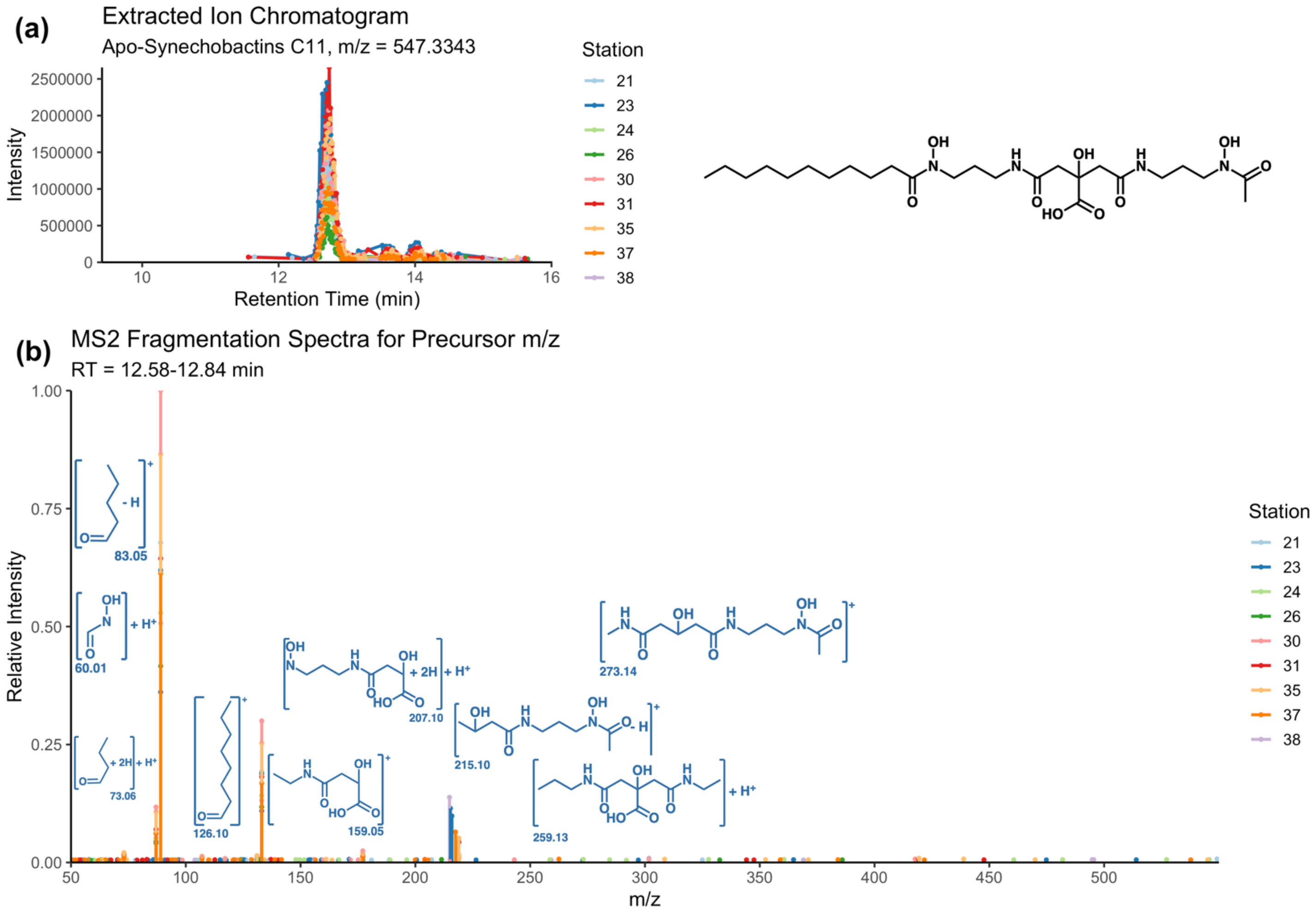

**Fig. S7. Confidence Level 2 example MS$^1$ and MS$^2$ spectra.** In a similar format to **Fig. S6**, example MS$^1$ (a) and MS$^2$ (b) spectra of putative siderophore Apo-Synechobactins $C_{11}$ is presented as an example of putative siderophores identified at confidence level 2. The peak in the extracted ion chromatogram (EIC) corresponding to the apo mass is presented in (a). Since this candidate structure is the apo form and not the *m/z* bound to Fe, the $^{54}$Fe isotopologue EIC is not plotted. Following analysis of the EIC, MS$^2$ fragments were extracted from the same *m/z* and retention time window and compared against *in silico* spectra for the candidate structure generated by MetFrag. Fragments found in MS$^2$ spectra that were also predicted by MetFrag are shown in (b). Many of these fragment masses were low-intensity in the MS$^2$ data, except for the fragment *m/z* 215.10 detected at St. 38, which was one of the most intense fragments in that sample. The well-defined MS$^1$ peaks and siderophore-like fragments matching *in silico* fragmentation — albeit at lower relative intensities — yields a detection at confidence level 2 for most sites. The detection at St. 38, however, was assigned a confidence level 1 due to the fragment with *m/z* = 215.10.

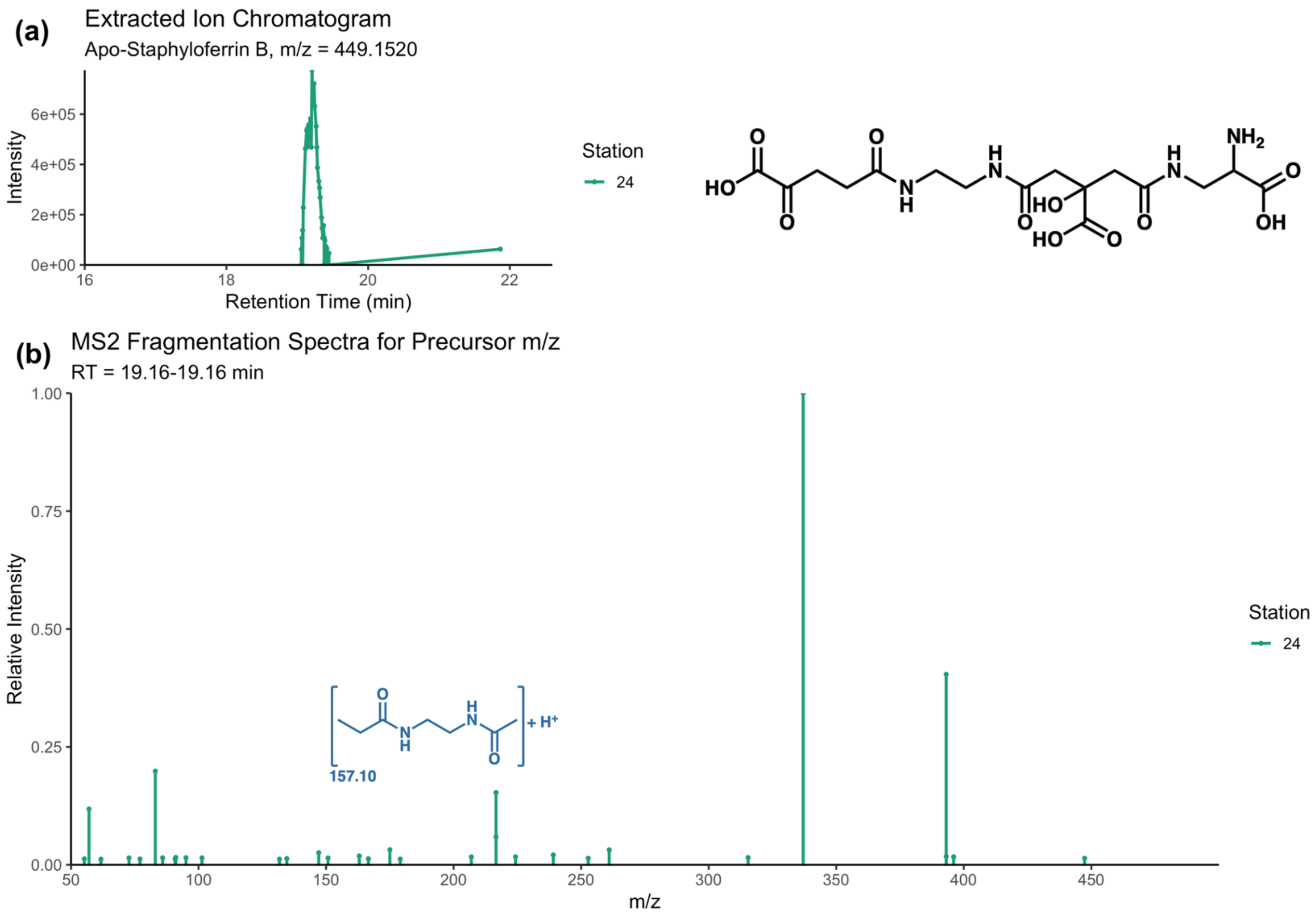

**Fig. S8. Confidence Level 3 example $MS^1$ and $MS^2$ spectra.** In a similar format to **Fig. S6**, example $MS^1$ (a) and $MS^2$ (b) spectra of putative siderophore Apo-Staphyloferrin B is presented as an example of putative siderophores identified at confidence level 3. The peak in the extracted ion chromatogram (EIC) corresponding to the apo mass is presented in (a). Since this candidate structure is the apo form and not the *m/z* bound to Fe, the $^{54}Fe$ isotopologue EIC is not plotted. Following analysis of the EIC, $MS^2$ fragments were extracted from the same *m/z* and retention time window and compared against *in silico* spectra for the candidate structure generated by MetFrag. Only one fragment was found in $MS^2$ spectra that was also predicted by MetFrag (b). While there was a well-defined $MS^1$ peaks that corresponded to the exact mass of Staphyloferrin B, only one fragment matched *in silico* fragmentation, yielding a detection at confidence level 3 at St. 24. These low-confidence detections are not included in most of the figures and relationships presented in the main text (unless detections were made at higher confidence at other sites and retention times align) but are presented in the supplement to aid future work.

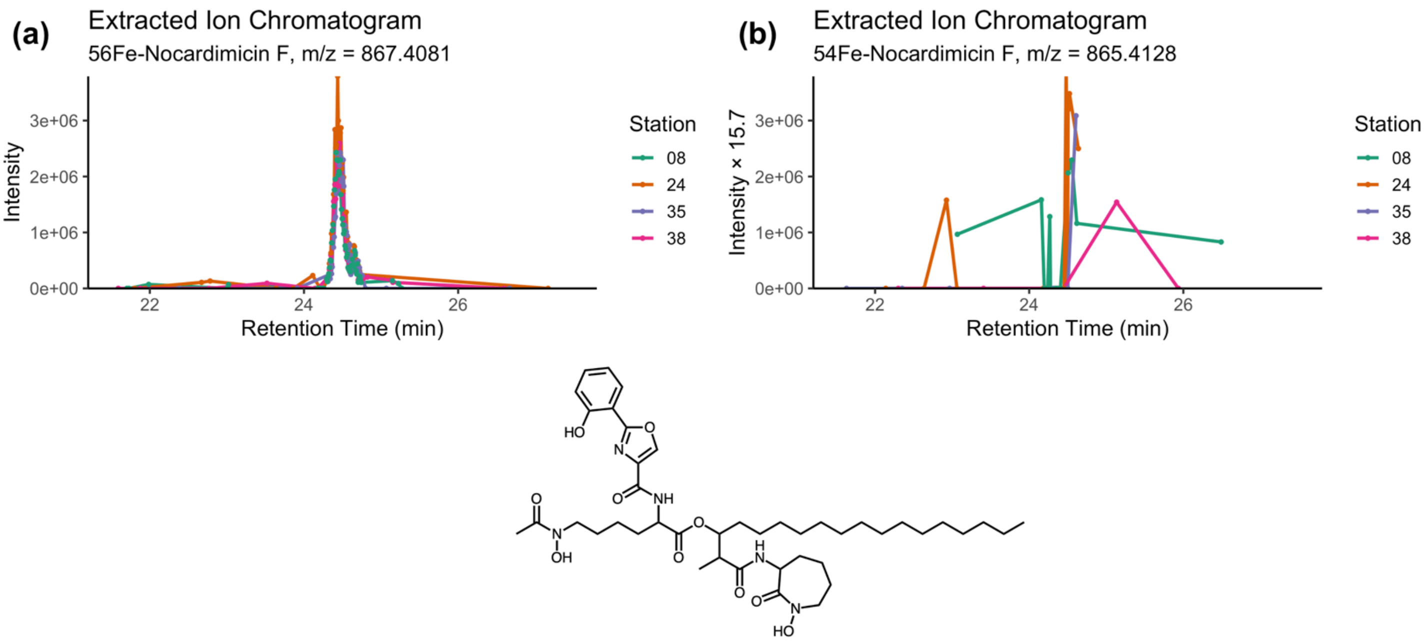

**Fig. S9. Confidence Level 4 example $MS^1$ spectra.** In a similar format to **Fig. S6**, example $MS^1$ (a-b) spectra of putative siderophore Fe-Nocardimicin F is presented as an example of putative siderophores identified at the lowest confidence level, 4. The peak in the extracted ion chromatogram (EIC) corresponding the $^{56}$Fe-bound mass is presented in (a), while the EIC corresponding to the less-abundant $^{54}$Fe-bound mass is presented in (b), scaled by a factor of 15.7 (crustal abundance of $^{56}$Fe/$^{54}$Fe). Little is known about the isotopic fractionation associated with Fe-siderophore binding, but the fact that the $^{54}$Fe-Nocardimicin F signal scales to a similar intensity to that of $^{56}$Fe-Nocardimicin F provides strong evidence that the candidate compound is indeed bound to Fe. That said, no $MS^2$ spectra was available for this candidate *m/z*, yielding a confidence level of 4. These low-confidence detections are not included in most of the figures and relationships presented in the main text (unless detections were made at higher confidence at other sites and retention times align) but are presented in the supplement to aid future work. Interestingly, Nocardimicin F and Terpenibactin A are very structurally similar and the $MS^1$ peaks in our data corresponding to their masses were of similar magnitude. The *m/z* of Nocardimicin F is within the 7.5 ppm tolerance of the *m/z* Terpenibactin A minus two hydrogen atoms, so it is possible that this compound is related to the putative Terpenibactin A compound through one degree of unsaturation in the hydrocarbon chain, which is common in marine siderophores (3). Our data, however, is unable to resolve this level of structural detail.

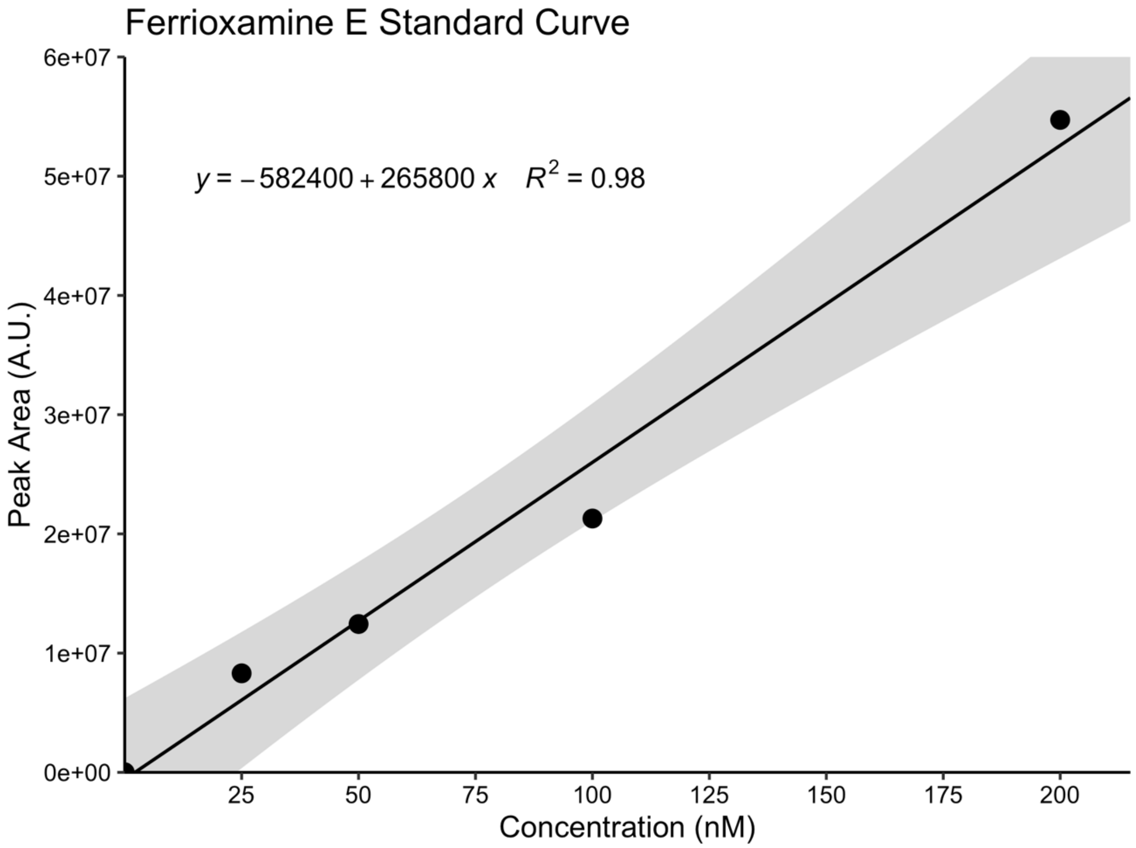

**Fig. S10. Ferrioxamine E Standard Curve.** A 5-point standard curve with known concentrations of siderophore ferrioxamine E at 0 nM, 25 nM, 50nM, 100 nM, and 200nM was used for quantification of putative siderophores. Commercial standards are not available for most siderophores, and different compounds have distinct ionization efficiencies in ESI-MS. Thus, the siderophore concentrations reported here are estimates of siderophore concentrations in these environments based on the structure and ionization efficiency of siderophore ferrioxamine E. The standard error of the slope of the curve was 20726 a.u./nM, yielding a limit of detection in the eluent of 0.257 nM. Eluent concentrations of 0.257 nM equate to sample concentrations of 0.07-0.21 pM depending on sample-to-eluent volume ratio at each site (**Table S1**). All putative detections below this limit were ultimately discarded. Additionally, 1 mM of cyanocobalamin was added as an internal standard to each sample aliquot to address any changes in sensitivity during LC-ESI-MS runs. Shaded area in the figure represents the 95% confidence interval of the linear regression.

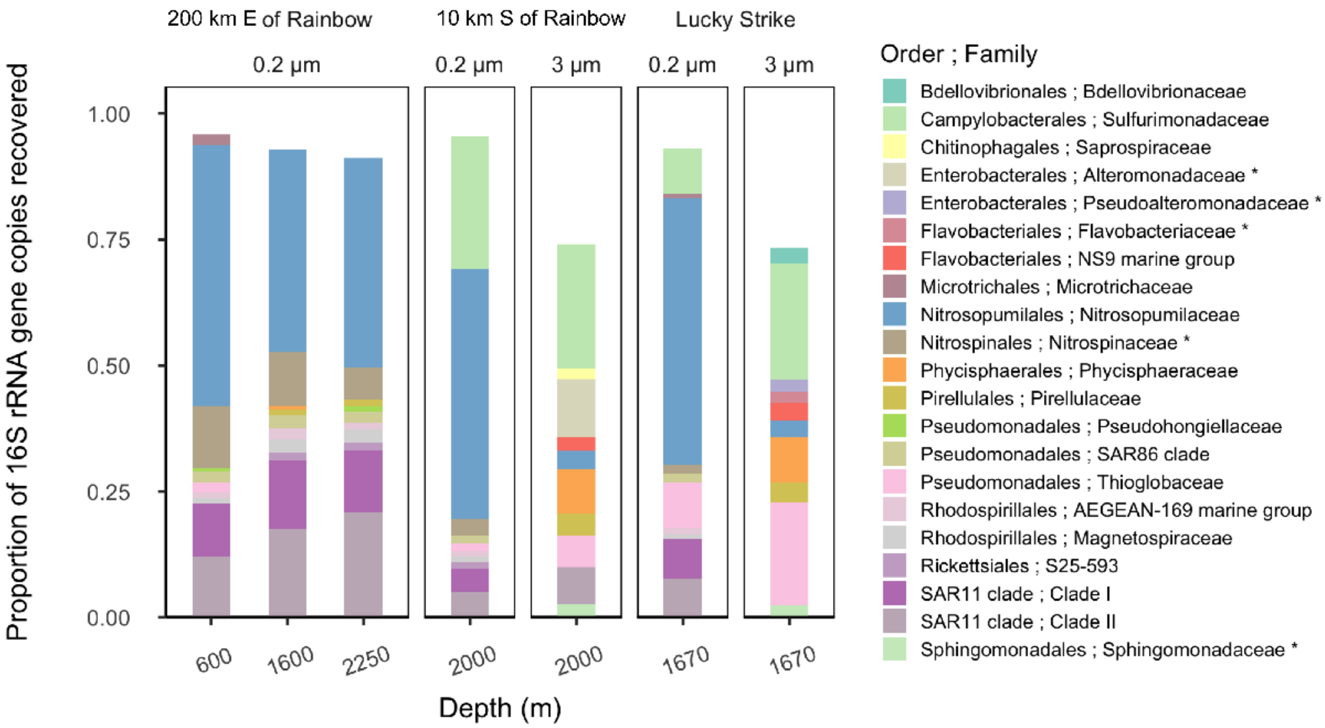

**Fig. S11. Relative abundance of prokaryotic taxa.** Bar height indicates the proportion of 16S rRNA genes recovered in each sample, separated by depth and site location. Colors correspond to taxonomy. The 10 most abundant families in each sample are depicted. Asterisks denote families containing genera hypothesized to produce siderophores.

**S3. Supporting Information Tables**

**Table S1. Fridge sample processing information**

| Location | Abbr. | Station # | Total # of Samples | # of CSV samples | 16S rRNA | Volume of seawater on SPE column (L)* | Volume of Eluent (μL)* |
|---|---|---|---|---|---|---|---|
| Lucky Strike† | LS | 7 | - | - | yes | - | - |
| Lucky Strike† | LS | 8 | 6 | 3 | | 1.50 | 416.1 |
| 200 km E of Rainbow | - | 11 | - | - | yes | - | - |
| 33 km E of Rainbow | CER | 12 | 5 | 2 | - | 0.95 | 500 |
| Rainbow | R | 38 | 3 | 1 | - | 0.65 | 542.4 |
| 10 km S of Rainbow | - | 17 | - | - | yes | - | - |
| Hayes Fracture Zone | HFZ | 21 | 6 | 3 | - | 1.05 | 500 |
| Lost City | LC | 23 | 8 | 6 | - | 1.10 | 500 |
| Broken Spur | BS | 24 | 8 | 4 (2) | - | 1.20 | 401.2 |
| 29 km N of TAG | CNT | 26 | 7 | 5 | - | 1.00 | 500 |
| 30 km W of TAG | CWT | 30 | 6 | 3 | - | 1.05 | 500 |
| 30 km E of TAG | CET | 31 | 7 | 5 | - | 1.00 | 500 |
| TAG | TAG | 35 | 10 | 2 (8) | - | 1.40 | 434.2 |
| Low Temp Slope | LTS | 37 | 8 | 6 | - | 1.10 | 500 |

For CSV, 150 mL and 100 mL of water was needed to run forward titrations and reverse titrations (in paratheses), respectively.

*For LC-MS, water samples were pooled and filtered from each vent site – see methods section for more details. Both volume of seawater concentrated, and final eluent volume are reported above.

- = not applicable/not sampled

† = Lucky Strike Station 7 and 8 were sampled at the same latitude and longitude (**Fig. 1a**)

**Table S2. Ligand concentrations and binding strengths from samples taken along the MAR.**

| Vent Name | Station | GT # | Depth<br>m | [dFe]<br>nM | $^{3}He_{xs}$-†<br>fmol | [$L_1$]<br>nM | [$L_2$]<br>nM | [$L_3$]<br>nM | [$L_T$]<br>nM | Log $K_1$ | Log $K_2$ | Log $K_3$ |
|---|---|---|---|---|---|---|---|---|---|---|---|---|
| BS | 24 | 1251 | 2950 | 0.94 | *n.a.* | | 1.62 ± 0.2 | | 1.62 ± 0.2 | | 11.922 ± 0.387 | |
| BS | 24 | 1252 | 2887 | 2.75 | *n.a.* | 3.21 ± 0.06 | | 37.1 ± 3.2 | 40.31 ± 3.26 | 13.42 ± 0.114 | | 9.778 ± 0.044 |
| BS | 24 | 1253 | 2841 | 7.57 | *n.a.* | 8.25 ± 0.47 | | | 8.25 ± 0.47 | 12.131 ± 0.364 | | |
| BS | 24 | 1255* | 2833 | 21.31 | *n.a.* | | 4.24±0.355 | | 4.24±0.355 | | 11.7±0.3 | |
| BS | 24 | 1258* | 2809 | 21.06 | *n.a.* | | 6.31±0.664 | | 6.31±0.664 | | 11.82±0.417 | |
| BS | 24 | 1260 | 2500 | 1.7 | *n.a.* | 2.17 ± 0.28 | | 58.5 ± 26.3 | 60.67 ± 26.58 | 12.434 ± 0.466 | | 8.745 ± 0.219 |
| CER | 12 | 628 | 2200 | 2.6 | 0.323 | 3.45 ± 0.43 | | 10.4 ± 4.0 | 13.85 ± 4.43 | 12.091 ± 0.363 | | 9.811 ± 0.221 |
| CER | 12 | 632 | 2000 | 2.61 | 0.313 | 3.98 ± 0.28 | | | 3.98 ± 0.28 | 12.132 ± 0.273 | | |
| CET | 31 | 1610 | 3700 | 0.49 | 0.191 | | | 4.93±0.89 | 4.93±0.89 | | | 10.509±0.158 |
| CET | 31 | 1612 | 3350 | 0.41 | 0.191 | | 2.39 ± 0.31 | 69.4 ± 30.4 | 71.79 ± 30.71 | | 11.469 ± 0.172 | 8.75 ± 0.219 |
| CET | 31 | 1613 | 3150 | 0.47 | 0.191 | | 3.65 ± 0.365 | | 3.65 ± 0.365 | | 11.054 ± 0.138 | |
| CET | 31 | 1615 | 2800 | 0.64 | 0.191 | 0.714 ± 0.143 | | 79.7 ± 24.9 | 80.414 ± 25.043 | 12.601 ± 0.662 | | 8.837 ± 0.155 |
| CET | 31 | 1616 | 2600 | 0.8 | 0.192 | | 2.25±0.27 | | 2.25±0.27 | | 11.504±0.201 | |
| CNT | 26 | 1346 | 3801 | 1.93 | 0.205 | | 5.57 ± 0.45 | 13.4 ± 5.3 | 18.97 ± 5.75 | | 11.748 ± 0.117 | 9.677 ± 0.242 |
| CNT | 26 | 1348 | 3400 | 3.87 | 0.232 | 6.96 ± 0.28 | | | 6.96 ± 0.28 | 12.06 ± 0.096 | | |
| CNT | 26 | 1349 | 3200 | 2.86 | 0.230 | | | 5.64 ± 0.99 | 5.64 ± 0.99 | | | 10.583 ± 0.238 |

| | | | | | | | | | | | | |
|---|---|---|---|---|---|---|---|---|---|---|---|---|
| CNT | 26 | 1351 | 2800 | 1.62 | 0.202 | 2.4 ± 0.16 | | | 2.4 ± 0.16 | 12.018 ± 0.21 | | |
| CNT | 26 | 1352 | 2400 | 0.89 | 0.198 | 1.22 ± 0.05 | | 2.05 ± 0.22 | 3.27 ± 0.27 | 12.778 ± 0.415 | | 10.663 ± 0.075 |
| CWT | 30 | 1539 | 2800 | 0.87 | 0.193 | | 1.6 ± 0.2 | | 1.6 ± 0.2 | | 11.636 ± 0.291 | |
| CWT | 30 | 1540 | 2700 | 1.22 | 0.191 | 1.91 ± 0.07 | | 25.3 ± 6.4 | 27.21 ± 6.47 | 12.547 ± 0.22 | | 9.026 ± 0.158 |
| CWT | 30 | 1542 | 2500 | 0.82 | 0.192 | | 2.77 ± 0.37 | | 2.77 ± 0.37 | | 11.228 ± 0.168 | |
| HFZ | 21 | 1129 | 4452 | 0.74 | *n.a.* | 0.697 ± 0.139 | | 34.9 ± 6.6 | 35.597 ± 6.739 | 12.419 ± 0.528 | | 9.515 ± 0.109 |
| HFZ | 21 | 1136 | 2850 | 1.31 | 0.161 | 1.38 ± 0.07 | | 10.2 ± 3.2 | 11.58 ± 3.27 | 13.189 ± 0.396 | | 9.592 ± 0.168 |
| HFZ | 21 | 1138 | 2500 | 1.084 | 0.163 | | 1.85 ± 0.08 | 5.58 ± 2.0 | 7.43 ± 2.08 | | 11.998 ± 0.108 | 9.19 ± 0.185 |
| LC | 23 | 1203 | 757 | 0.63 | *n.a.* | 1.24 ± 0.16 | | 9.05 ± 3.06 | 10.29 ± 3.22 | 12.23 ± 0.336 | | 9.676 ± 0.194 |
| LC | 23 | 1204 | 753 | 0.81 | *n.a.* | | 6.19 ± 0.34 | | 6.19 ± 0.34 | | 11.164 ± 0.067 | |
| LC | 23 | 1205 | 747 | 0.73 | *n.a.* | 1.238 ± 0.062 | | 8.086 ± 1.33 | 9.324 ± 1.392 | 12.02 ± 0.114 | | 9.573 ± 0.081 |
| LC | 23 | 1207 | 727 | 0.51 | *n.a.* | 1.005 ± 0.075 | | 2.244 ± 0.269 | 3.249 ± 0.344 | 12.115 ± 0.151 | | 10.645 ± 0.09 |
| LC | 23 | 1208 | 700 | 0.78 | *n.a.* | 1.27 ± 0.08 | | 15 ± 3.5 | 16.27 ± 3.58 | 12.189 ± 0.171 | | 9.319 ± 0.14 |
| LC | 23 | 1209 | 685 | 0.58 | *n.a.* | | 2.527 ± 0.215 | | 2.527 ± 0.215 | | 11.346 ± 0.091 | |
| LTS | 37 | 1970 | 2400 | 0.9 | 0.192 | 1.43 ± 0.2 | | 24.2 ± 11.2 | 25.63 ± 11.4 | 12.151 ± 0.365 | | 9.163 ± 0.229 |
| LTS | 37 | 1972 | 2200 | 0.87 | 0.192 | | 1.78 ± 0.19 | 7.08 ± 2.04 | 8.86 ± 2.23 | | 11.531 ± 0.173 | 9.624 ± 0.168 |
| LTS | 37 | 1973 | 2100 | 0.73 | 0.191 | 1.3 ± 0.07 | | | 1.3 ± 0.07 | 12.191 ± 0.244 | | |
| LTS | 37 | 1975 | 1900 | 0.71 | 0.191 | | 1.47 ± 0.24 | | 1.47 ± 0.24 | | 11.789 ± 0.295 | |

| | | | | | | | | | | | | |
|---|---|---|---|---|---|---|---|---|---|---|---|---|
| LTS | 37 | 1976 | 1800 | 0.65 | 0.191 | 1.3 ± 0.06 | | | 1.3 ± 0.06 | 12.048 ± 0.151 | | |
| LTS | 37 | 1977 | 1700 | 0.59 | 0.191 | 1.38 ± 0.14 | | | 1.38 ± 0.14 | 12.179 ± 0.365 | | |
| LS | 8 | 409 | 1725 | 11.56 | 0.902 | 12.8 ± 0.3 | | | 12.8 ± 0.3 | 13.096 ± 0.458 | | |
| LS | 8 | 415 | 1620 | 11.67 | 0.840 | | 12.6 ± 0.3 | | 12.6 ± 0.3 | | 11.924±0.143 | |
| LS | 8 | 418 | 1550 | 1.28 | 0.505 | 2.34 ± 0.13 | | | 2.34 ± 0.13 | 12.519 ± 0.72 | | |
| R | 38 | 1009 | 2285 | 25.87 | 0.446 | | | 27.3 ± 1.3 | 27.3 ± 1.3 | | | 10.967 ± 0.192 |
| TAG | 35 | 1802 | 3600 | 2.9 | 0.212 | 3.59 ± 0.22 | | | 3.59 ± 0.22 | 12.022 ± 0.601 | | |
| TAG | 35 | 1806* | 3453 | 19.01 | 0.284 | | | 9.19±0.538 | 9.19±0.538 | | | 10.66±0.166 |
| TAG | 35 | 1807* | 3429 | 26.99 | 0.326 | | | 8.27±0.539 | 8.27±0.539 | | | 10.68±0.178 |
| TAG | 35 | 1811* | 3335 | 30.68 | 1.122 | | | 20.1±2.66 | 20.1±2.66 | | | 10.22±0.361 |
| TAG | 35 | 1812* | 3322 | 64.36 | 0.788 | | | 6.14±1.64 | 6.14±1.64 | | | 10.58±0.677 |
| TAG | 35 | 1813* | 3313 | 34.22 | 0.777 | | | | | | | |
| TAG | 35 | 1817* | 3336 | 90.25 | 0.849 | | | 37.7±5.93 | 37.7±5.93 | | | 10.04±0.438 |
| TAG | 35 | 1818* | 3195 | 84.53 | 0.431 | | 9.55±1.59 | | 9.55±1.59 | | 11.58±0.654 | |
| TAG | 35 | 1819* | 3100 | 55.71 | 0.201 | | | | | | | |
| TAG | 35 | 1821 | 1821 | 1.34 | 0.192 | | 2.26 ± 0.35 | | 2.26 ± 0.35 | | 11.689 ± 0.497 | |

$L_1$, $L_2$, and $L_3$ are defined in the literature(4). Reverse titrations (*) were not found to contain $L_1$ ligands. GT# 1813 and 1819 titrations were not successful.

*reverse titration

† Expected $^3He_{xs}$ values were derived from the conservative relationship between dMn/$^3He_{xs}$ in hydrothermal plumes(1).

*n.a.* = $^3He_{xs}$ data not available

**Table S3. Average Strong binding ligands and Siderophore concentrations along the MAR**

| Location | Station | Avg $L_1$ ligand (nM) | Avg Siderophore Concentration (pM) |
|---|---|---|---|
| Lucky Strike | 8 | 7.57±7.40 | 0.438±0.042 |
| 33 km E of Rainbow | 12 | 3.72±0.37 | 0.269±0.080 |
| Hayes Fracture Zone | 21 | 1.04±0.48 | 0.548±0.073 |
| Lost City | 23 | 1.24±0.15 | 0.579±0.069 |
| Broken Spur | 24 | 4.54±3.25 | 0.450±0.051 |
| 29 km N of TAG | 26 | 3.53±3.03 | 0.242±0.076 |
| 30 km W of TAG | 30 | 1.91 | 0.969±0.073 |
| 30 km E of TAG | 31 | 0.71 | 0.548±0.076 |
| TAG | 35 | 3.59 | 0.440±0.047 |
| Low Temp Slope | 37 | 1.35±0.06 | 0.282±0.069 |
| Rainbow | 38 | *n.a.* | 0.863±0.128 |

Standard deviations are reported for average $L_1$ concentrations, while, for average siderophore concentrations, the error is reported as 1.96*SE of the desferrioxamine E standard curve multiplied by the concentration factor of each sample.

*n.a.* = no $L_1$ ligands were found in the sample(s) that were run using CSV.

**Table S4. Brief description of confidence levels**

| Confidence Level | $MS^1$ | $MS^2$ | Fragmentation Pattern |
|---|---|---|---|
| 1* | ✓ | ✓ | Multiple fragments in the $MS^2$ data matched those predicted by MetFrag and at least one was in the top three most intense fragments |
| 2* | ✓ | ✓ | Multiple lower-intensity fragments in the $MS^2$ data matched those predicted by MetFrag |
| 3 | ✓ | ✓ | Little to no fragments found in $MS^2$ data match those predicted by MetFrag |
| 4 | ✓ | - | - |

*Confidence levels were designed such that putative detections made with either confidence level 1 or 2 contained siderophore-like fragments in $MS^2$ data, regardless of definitive identification. That is, we have reasonable confidence that the putative siderophore at least contains siderophore moieties.

**Table S5. Putative siderophores detected with LC-ESI-MS**

| Vent Name | Station | Geology | Putative Siderophore | Form | *m/z* | Confidence Level | Type | Hydrocarbon Chain?* | Retention Time (min) | Raw Peak Area | Peak Area × Conc. Factor | DFOE-calibrated Conc. in SW (pM) | Conc. in SW with 40% Efficiency (pM)** | Conc. in SW with 10% Efficiency (pM)*** |
|---|---|---|---|---|---|---|---|---|---|---|---|---|---|---|
| LS | 8 | Spreading Center | Terpenibactin A | Fe | 869.424 | 1 | Mixed | Yes | 24.8 | 377950 | 104.843 | 0.394 | 0.986 | 3.944 |
| LS | 8 | Spreading Center | Acremonpeptide B | Apo | 848.452 | 1 | Hydroxamate | No | 24.8 | 476264 | 132.115 | 0.497 | 1.243 | 4.970 |
| LS | 8 | Spreading Center | Mycobactin S/T C17 | Apo | 828.512 | 1 | Mixed | Yes | 15 | 180723 | 50.133 | 0.189 | 0.471 | 1.886 |
| CER | 12 | Spreading Center | Mycobactin S/T C17 | Apo | 828.512 | 1 | Mixed | Yes | 15 | 151918 | 79.957 | 0.301 | 0.752 | 3.008 |
| HFZ | 21 | Fracture Zone | Mycobactin S/T C17 | Apo | 828.512 | 1 | Mixed | Yes | 15 | 507896 | 241.855 | 0.910 | 2.275 | 9.098 |
| LC | 23 | Fracture Zone | Mycobactin S/T C17 | Apo | 828.512 | 1 | Mixed | Yes | 15 | 707028 | 321.376 | 1.209 | 3.023 | 12.090 |
| LC | 23 | Fracture Zone | Woodybactin D | Fe | 544.172 | 1 | Carboxylate | Yes | 16.9 | 103258 | 46.936 | 0.177 | 0.441 | 1.766 |
| LC | 23 | Fracture Zone | Thiazostatin | Apo | 339.084 | 1 | Mixed | No | 19 | 87227 | 39.649 | 0.149 | 0.373 | 1.492 |
| BS | 24 | Spreading Center | Amphibactin D | Fe | 885.415 | 1 | Hydroxamate | Yes | 24.8 | 353010 | 118.023 | 0.444 | 1.110 | 4.440 |
| BS | 24 | Spreading Center | Nocardimicin I +H | Fe | 884.447 | 1 | Mixed | Yes | 24.5 | 159387 | 53.289 | 0.200 | 0.501 | 2.005 |
| BS | 24 | Spreading Center | Terpenibactin A | Fe | 869.424 | 1 | Mixed | Yes | 24.8 | 577343 | 193.025 | 0.726 | 1.815 | 7.262 |
| BS | 24 | Spreading Center | Acremonpeptide B | Apo | 848.452 | 1 | Hydroxamate | No | 24.8 | 790407 | 264.259 | 0.994 | 2.485 | 9.941 |
| BS | 24 | Spreading Center | Mycobactin S/T C17 | Apo | 828.512 | 1 | Mixed | Yes | 15 | 296850 | 99.247 | 0.373 | 0.933 | 3.734 |
| CNT | 26 | Spreading Center | Mycobactin S/T C17 | Apo | 828.512 | 1 | Mixed | Yes | 15 | 213887 | 106.943 | 0.402 | 1.006 | 4.023 |
| CWT | 30 | Spreading Center | Mycobactin S/T C17 | Apo | 828.512 | 1 | Mixed | Yes | 15 | 509863 | 242.792 | 0.913 | 2.283 | 9.134 |

| | | | | | | | | | | | | | | |
|---|---|---|---|---|---|---|---|---|---|---|---|---|---|---|
| CWT | 30 | Spreading Center | Woodybactin D | Fe | 544.172 | 1 | Carboxylate | Yes | 16.9 | 770593 | 366.949 | 1.380 | 3.451 | 13.804 |
| CET | 31 | Spreading Center | Mycobactin S/T C17 | Apo | 828.512 | 1 | Mixed | Yes | 15 | 556283 | 278.142 | 1.046 | 2.616 | 10.464 |
| CET | 31 | Spreading Center | Woodybactin D | Fe | 544.172 | 1 | Carboxylate | Yes | 16.9 | 90528 | 45.264 | 0.170 | 0.426 | 1.703 |
| TAG | 35 | Spreading Center | Nocardimicin I +H | Fe | 884.447 | 1 | Mixed | Yes | 24.5 | 101851 | 31.588 | 0.119 | 0.297 | 1.188 |
| TAG | 35 | Spreading Center | Terpenibactin A | Fe | 869.424 | 1 | Mixed | Yes | 24.8 | 414993 | 128.707 | 0.484 | 1.210 | 4.842 |
| TAG | 35 | Spreading Center | Acremonpeptide B | Apo | 848.452 | 1 | Hydroxamate | No | 24.8 | 508219 | 157.621 | 0.593 | 1.482 | 5.930 |
| TAG | 35 | Spreading Center | Mycobactin S/T C17 | Apo | 828.512 | 1 | Mixed | Yes | 15 | 491344 | 152.387 | 0.573 | 1.433 | 5.733 |
| TAG | 35 | Spreading Center | Woodybactin D | Fe | 544.172 | 1 | Carboxylate | Yes | 16.9 | 81066 | 25.142 | 0.095 | 0.236 | 0.946 |
| LTS | 37 | Fracture Zone | Mycobactin S/T C17 | Apo | 828.512 | 1 | Mixed | Yes | 15 | 219752 | 99.887 | 0.376 | 0.939 | 3.758 |
| R2 | 38 | Spreading Center | Nocardimicin I +H | Fe | 884.447 | 1 | Mixed | Yes | 24.5 | 89956 | 75.065 | 0.282 | 0.706 | 2.824 |
| R2 | 38 | Spreading Center | Terpenibactin A | Fe | 869.424 | 1 | Mixed | Yes | 24.8 | 358436 | 299.101 | 1.125 | 2.813 | 11.252 |
| R2 | 38 | Spreading Center | Acremonpeptide B | Apo | 848.452 | 1 | Hydroxamate | No | 24.8 | 472652 | 394.410 | 1.484 | 3.709 | 14.838 |
| R2 | 38 | Spreading Center | Mycobactin S/T C17 | Apo | 828.512 | 1 | Mixed | Yes | 15 | 426981 | 356.299 | 1.340 | 3.351 | 13.404 |
| R2 | 38 | Spreading Center | Synechobactins C11 | Apo | 547.334 | 1 | Mixed | Yes | 12.7 | 261667 | 218.351 | 0.821 | 2.054 | 8.214 |
| LS | 8 | Spreading Center | Amphibactin D | Fe | 885.415 | 2 | Hydroxamate | Yes | 24.7 | 229587 | 63.688 | 0.240 | 0.599 | 2.396 |
| LS | 8 | Spreading Center | IC202C | Fe | 570.246 | 2 | Hydroxamate | No | 23.5 | 363338 | 100.790 | 0.379 | 0.948 | 3.792 |
| LS | 8 | Spreading Center | Chlorocatechelin B | Apo | 461.144 | 2 | Mixed | No | 18.9 | 428902 | 118.977 | 0.448 | 1.119 | 4.476 |
| LS | 8 | Spreading Center | Vibrioferrin C-2H | Apo | 447.125 | 2 | Carboxylate | No | 14.7 | 581310 | 161.255 | 0.607 | 1.517 | 6.066 |

| | | | | | | | | | | | | | | |
|---|---|---|---|---|---|---|---|---|---|---|---|---|---|---|
| HFZ | 21 | Fracture Zone | Synechobactins C11 | Apo | 547.334 | 2 | Mixed | Yes | 12.7 | 282675 | 134.607 | 0.506 | 1.266 | 5.064 |
| HFZ | 21 | Fracture Zone | Woodybactin D | Fe | 544.172 | 2 | Carboxylate | Yes | 16.9 | 147103 | 70.049 | 0.264 | 0.659 | 2.635 |
| HFZ | 21 | Fracture Zone | Vibrioferrin C-2H | Apo | 447.125 | 2 | Carboxylate | No | 14.7 | 422356 | 201.122 | 0.757 | 1.892 | 7.566 |
| LC | 23 | Fracture Zone | Acinetoferrin | Apo | 585.350 | 2 | Mixed | Yes | 18.6 | 101336 | 46.062 | 0.173 | 0.433 | 1.733 |
| LC | 23 | Fracture Zone | Synechobactins C11 | Apo | 547.334 | 2 | Mixed | Yes | 12.7 | 429185 | 195.084 | 0.734 | 1.835 | 7.339 |
| LC | 23 | Fracture Zone | Vibrioferrin C-2H | Apo | 447.125 | 2 | Carboxylate | No | 14.7 | 602509 | 273.868 | 1.030 | 2.576 | 10.303 |
| BS | 24 | Spreading Center | Crochelin A | Apo | 696.353 | 2 | Mixed | No | 26 | 377773 | 126.302 | 0.475 | 1.188 | 4.751 |
| BS | 24 | Spreading Center | IC202C | Fe | 570.246 | 2 | Hydroxamate | No | 23.5 | 534621 | 178.742 | 0.672 | 1.681 | 6.724 |
| BS | 24 | Spreading Center | Synechobactins C11 | Apo | 547.334 | 2 | Mixed | Yes | 12.7 | 164152 | 54.882 | 0.206 | 0.516 | 2.065 |
| BS | 24 | Spreading Center | Oxahomorhizoferrin | Fe | 506.047 | 2 | Carboxylate | No | 23.4 | 290782 | 97.218 | 0.366 | 0.914 | 3.657 |
| BS | 24 | Spreading Center | Chlorocatechelin B | Apo | 461.144 | 2 | Mixed | No | 18.9 | 75889 | 25.372 | 0.095 | 0.239 | 0.954 |
| BS | 24 | Spreading Center | Vibrioferrin C-2H | Apo | 447.125 | 2 | Carboxylate | No | 14.7 | 465962 | 155.787 | 0.586 | 1.465 | 5.861 |
| CNT | 26 | Spreading Center | Synechobactins C11 | Apo | 547.334 | 2 | Mixed | Yes | 12.7 | 88291 | 44.145 | 0.166 | 0.415 | 1.661 |
| CNT | 26 | Spreading Center | Chlorocatechelin B | Apo | 461.144 | 2 | Mixed | No | 18.9 | 101696 | 50.848 | 0.191 | 0.478 | 1.913 |
| CWT | 30 | Spreading Center | Synechobactins C11 | Apo | 547.334 | 2 | Mixed | Yes | 12.7 | 341539 | 162.638 | 0.612 | 1.530 | 6.118 |
| CET | 31 | Spreading Center | Synechobactins C11 | Apo | 547.334 | 2 | Mixed | Yes | 12.7 | 420695 | 210.347 | 0.791 | 1.978 | 7.913 |
| TAG | 35 | Spreading Center | Amphibactin D | Fe | 885.415 | 2 | Hydroxamate | Yes | 24.6 | 257644 | 79.907 | 0.301 | 0.752 | 3.006 |
| TAG | 35 | Spreading Center | Petrobactin Sulfonate | Apo | 771.287 | 2 | Mixed | No | 24.2 | 1046933 | 324.699 | 1.222 | 3.054 | 12.215 |

| | | | | | | | | | | | | | | |
|---|---|---|---|---|---|---|---|---|---|---|---|---|---|---|
| TAG | 35 | Spreading Center | Desferrioxamine P1 | Apo | 616.331 | 2 | Hydroxamate | No | 26.5 | 791716 | 245.545 | 0.924 | 2.309 | 9.237 |
| TAG | 35 | Spreading Center | Acinetoferrin | Apo | 585.350 | 2 | Mixed | Yes | 18.6 | 159590 | 49.496 | 0.186 | 0.466 | 1.862 |
| TAG | 35 | Spreading Center | IC202C | Fe | 570.246 | 2 | Hydroxamate | No | 23.5 | 424015 | 131.505 | 0.495 | 1.237 | 4.947 |
| TAG | 35 | Spreading Center | Synechobactins C11 | Apo | 547.334 | 2 | Mixed | Yes | 12.8 | 374766 | 116.231 | 0.437 | 1.093 | 4.373 |
| TAG | 35 | Spreading Center | Oxahomorhizoferrin | Fe | 506.047 | 2 | Carboxylate | No | 23.4 | 381526 | 118.328 | 0.445 | 1.113 | 4.451 |
| TAG | 35 | Spreading Center | Chlorocatechelin B | Apo | 461.144 | 2 | Mixed | No | 18.9 | 206280 | 63.976 | 0.241 | 0.602 | 2.407 |
| TAG | 35 | Spreading Center | Vibrioferrin C-2H | Apo | 447.125 | 2 | Carboxylate | No | 14.7 | 343044 | 106.393 | 0.400 | 1.001 | 4.002 |
| LTS | 37 | Fracture Zone | Vibrioferrin C-2H | Apo | 447.125 | 2 | Carboxylate | No | 14.7 | 107803 | 49.001 | 0.184 | 0.461 | 1.843 |
| LTS | 37 | Fracture Zone | Fusarinine | Apo | 261.145 | 2 | Mixed | No | 16.6 | 145829 | 66.286 | 0.249 | 0.623 | 2.494 |
| R2 | 38 | Spreading Center | Amphibactin D | Fe | 885.415 | 2 | Hydroxamate | Yes | 24.7 | 232026 | 193.616 | 0.728 | 1.821 | 7.284 |
| R2 | 38 | Spreading Center | Acinetoferrin | Apo | 585.350 | 2 | Mixed | Yes | 18.6 | 117191 | 97.791 | 0.368 | 0.920 | 3.679 |
| R2 | 38 | Spreading Center | IC202C | Fe | 570.246 | 2 | Hydroxamate | No | 23.5 | 381777 | 318.578 | 1.198 | 2.996 | 11.985 |
| R2 | 38 | Spreading Center | Oxahomorhizoferrin | Fe | 506.047 | 2 | Carboxylate | No | 23.4 | 142918 | 119.259 | 0.449 | 1.122 | 4.486 |
| R2 | 38 | Spreading Center | Chlorocatechelin B | Apo | 461.144 | 2 | Mixed | No | 18.9 | 103894 | 86.696 | 0.326 | 0.815 | 3.261 |
| R2 | 38 | Spreading Center | Vibrioferrin C-2H | Apo | 447.125 | 2 | Carboxylate | No | 14.7 | 210102 | 175.322 | 0.660 | 1.649 | 6.596 |
| LS | 8 | Spreading Center | Petrobactin Sulfonate | Apo | 771.287 | 3 | Mixed | No | 24.1 | 480146 | 133.193 | 0.501 | 1.253 | 5.011 |
| LS | 8 | Spreading Center | Crochelin A | Apo | 696.353 | 3 | Mixed | No | 26 | 282119 | 78.260 | 0.294 | 0.736 | 2.944 |
| LS | 8 | Spreading Center | Desferrioxamine P1 | Apo | 616.331 | 3 | Hydroxamate | No | 26.5 | 797065 | 221.106 | 0.832 | 2.079 | 8.318 |

| | | | | | | | | | | | | | | |
|---|---|---|---|---|---|---|---|---|---|---|---|---|---|---|
| CER | 12 | Spreading Center | Desferrioxamine P1 | Apo | 616.331 | 3 | Hydroxamate | No | 26.5 | 119545 | 62.918 | 0.237 | 0.592 | 2.367 |
| BS | 24 | Spreading Center | Petrobactin Sulfonate | Apo | 771.287 | 3 | Mixed | No | 24.1 | 154561 | 51.675 | 0.194 | 0.486 | 1.944 |
| BS | 24 | Spreading Center | Desferrioxamine P1 | Apo | 616.331 | 3 | Hydroxamate | No | 26.5 | 632217 | 211.371 | 0.795 | 1.988 | 7.952 |
| BS | 24 | Spreading Center | Staphyloferrin B | Apo | 449.152 | 3 | Mixed | No | 19.3 | 132154 | 44.183 | 0.166 | 0.416 | 1.662 |
| R2 | 38 | Spreading Center | Crochelin A | Apo | 696.353 | 3 | Mixed | No | 26 | 206343 | 172.185 | 0.648 | 1.619 | 6.478 |
| R2 | 38 | Spreading Center | Desferrioxamine P1 | Apo | 616.331 | 3 | Hydroxamate | No | 26.5 | 768105 | 640.954 | 2.411 | 6.028 | 24.112 |
| LS | 8 | Spreading Center | Nocardimicin F | Fe | 867.408 | 4 | Mixed | Yes | 24.5 | 393157 | 109.062 | 0.410 | 1.026 | 4.103 |
| LS | 8 | Spreading Center | Desferrioxamine X4 | Fe | 682.299 | 4 | Hydroxamate | No | 28.2 | 126870 | 35.194 | 0.132 | 0.331 | 1.324 |
| HFZ | 21 | Fracture Zone | Fusarinine | Apo | 261.145 | 4 | Mixed | No | 16.5 | 169109 | 80.528 | 0.303 | 0.757 | 3.029 |
| LC | 23 | Fracture Zone | Mycobactin P C17 | Apo | 870.559 | 4 | Mixed | Yes | 21.4 | 72348 | 32.885 | 0.124 | 0.309 | 1.237 |
| LC | 23 | Fracture Zone | Nocardimicin I | Apo | 830.528 | 4 | Mixed | Yes | 15 | 80033 | 36.379 | 0.137 | 0.342 | 1.369 |
| LC | 23 | Fracture Zone | IC202A | Apo | 573.398 | 4 | Hydroxamate | No | 18.9 | 185180 | 84.173 | 0.317 | 0.792 | 3.167 |
| BS | 24 | Spreading Center | Nocardimicin F | Fe | 867.408 | 4 | Mixed | Yes | 24.5 | 523154 | 174.908 | 0.658 | 1.645 | 6.580 |
| BS | 24 | Spreading Center | Desferrioxamine T3 | Apo | 773.441 | 4 | Hydroxamate | No | 24.9 | 125850 | 42.076 | 0.158 | 0.396 | 1.583 |
| BS | 24 | Spreading Center | Desferrioxamine X4 | Fe | 682.299 | 4 | Hydroxamate | No | 28.2 | 130558 | 43.650 | 0.164 | 0.411 | 1.642 |
| BS | 24 | Spreading Center | Thiazostatin | Apo | 339.084 | 4 | Mixed | No | 19 | 136590 | 45.667 | 0.172 | 0.429 | 1.718 |
| CNT | 26 | Spreading Center | Talarazine A | Apo | 469.266 | 4 | Hydroxamate | No | 22 | 76585 | 38.293 | 0.144 | 0.360 | 1.441 |
| CNT | 26 | Spreading Center | Fusarinine | Apo | 261.145 | 4 | Mixed | No | 16.5 | 110730 | 55.365 | 0.208 | 0.521 | 2.083 |

| | | | | | | | | | | | | | | |
|---|---|---|---|---|---|---|---|---|---|---|---|---|---|---|
| CET | 31 | Spreading Center | Chlorocatechelin B | Apo | 461.144 | 4 | Mixed | No | 18.9 | 97801 | 48.901 | 0.184 | 0.460 | 1.840 |
| TAG | 35 | Spreading Center | Mycobactin P C17 | Apo | 870.559 | 4 | Mixed | Yes | 21.4 | 118442 | 36.734 | 0.138 | 0.345 | 1.382 |
| TAG | 35 | Spreading Center | Nocardimicin F | Fe | 867.408 | 4 | Mixed | Yes | 24.5 | 466429 | 144.660 | 0.544 | 1.361 | 5.442 |
| TAG | 35 | Spreading Center | Desferrioxamine T3 | Apo | 773.441 | 4 | Hydroxamate | No | 24.9 | 192467 | 59.692 | 0.225 | 0.561 | 2.246 |
| TAG | 35 | Spreading Center | Crochelin A | Apo | 696.353 | 4 | Mixed | No | 26 | 78347 | 24.299 | 0.091 | 0.229 | 0.914 |
| TAG | 35 | Spreading Center | Desferrioxamine X4 | Fe | 682.299 | 4 | Hydroxamate | No | 29 | 132006 | 40.941 | 0.154 | 0.385 | 1.540 |
| LTS | 37 | Fracture Zone | Synechobactins C11 | Apo | 547.334 | 4 | Mixed | Yes | 12.7 | 185568 | 84.349 | 0.317 | 0.793 | 3.173 |
| R2 | 38 | Spreading Center | Mycobactin P C17 | Apo | 870.559 | 4 | Mixed | Yes | 21.4 | 70466 | 58.801 | 0.221 | 0.553 | 2.212 |
| R2 | 38 | Spreading Center | Nocardimicin F | Fe | 867.408 | 4 | Mixed | Yes | 24.5 | 398350 | 332.408 | 1.251 | 3.126 | 12.505 |
| R2 | 38 | Spreading Center | Desferrioxamine X4 | Fe | 682.299 | 4 | Hydroxamate | No | 28.2 | 81880 | 68.326 | 0.257 | 0.643 | 2.570 |
| R2 | 38 | Spreading Center | Thiazostatin | Apo | 339.084 | 4 | Mixed | No | 19 | 78507 | 65.511 | 0.246 | 0.616 | 2.465 |

*A 'Yes' under 'Hydrocarbon Chain' indicates that the putative structure contains a hydrocarbon chain of 7 or more carbons

**40% efficiency is based on the efficiency of ENV columns for recovering siderophore desferrioxamine E in solid-phase extractions from seawater (5)

***10% efficiency is based on the typical recovery of the bulk Fe-binding organic pool with ENV columns, which comes from comparing the concentration of [Fe-L]-SPE from LC-ICP-MS with that of [Fe-L] from cathodic-stripping voltammetry (6)

**Table S6. Genera detected in this study along with corresponding siderophores production ability**

| Taxonomy | Relative Abundance | Location | Filter Size | Depth (m) | Putative Siderophores |
|---|---|---|---|---|---|
| Bacteria ; Nitrospinota ; Nitrospinia ; Nitrospinales ; Nitrospinaceae ; Nitrospina | 0.09466131 | 200 km E of Rainbow | 0.2 µm | 600 | STAPHYLOBACTIN |
| Bacteria ; Proteobacteria ; Gammaproteobacteria ; Enterobacterales ; Pseudoalteromonadaceae ; Pseudoalteromonas | 0.0013824 | 200 km E of Rainbow | 0.2 µm | 600 | DESFERRIOXAMINE E , PUTREBACTIN / AVAROFERRIN , DESFERRIOXAMIN B , DESFERRIOXAMINE , UNKNOWN , STAPHYLOBACTIN |
| Bacteria ; Proteobacteria ; Gammaproteobacteria ; Enterobacterales ; Colwelliaceae ; Colwellia | 0.00125074 | 200 km E of Rainbow | 0.2 µm | 600 | UNKNOWN |
| Bacteria ; Nitrospinota ; Nitrospinia ; Nitrospinales ; Nitrospinaceae ; Nitrospina | 0.09409973 | 200 km E of Rainbow | 0.2 µm | 1600 | STAPHYLOBACTIN |
| Bacteria ; Proteobacteria ; Gammaproteobacteria ; Enterobacterales ; Pseudoalteromonadaceae ; Pseudoalteromonas | 0.00350209 | 200 km E of Rainbow | 0.2 µm | 1600 | DESFERRIOXAMINE E , PUTREBACTIN / AVAROFERRIN , DESFERRIOXAMIN B , DESFERRIOXAMINE , UNKNOWN , STAPHYLOBACTIN |
| Bacteria ; Proteobacteria ; Gammaproteobacteria ; Pseudomonadales ; Marinobacteraceae ; Marinobacter | 0.00057099 | 200 km E of Rainbow | 0.2 µm | 1600 | PUTREBACTIN / AVAROFERRIN , XANTHOFERRIN , UNKNOWN |
| Bacteria ; Proteobacteria ; Gammaproteobacteria ; Enterobacterales ; Moritellaceae ; Moritella | 0.00030453 | 200 km E of Rainbow | 0.2 µm | 1600 | COLICIN V , UNKNOWN |
| Bacteria ; Proteobacteria ; Gammaproteobacteria ; Coxiellales ; Coxiellaceae ; Coxiella | 0.00019033 | 200 km E of Rainbow | 0.2 µm | 1600 | UNKNOWN |

| | | | | | |
|---|---|---|---|---|---|
| Bacteria ; Nitrospinota ; Nitrospinia ; Nitrospinales ; Nitrospinaceae ; Nitrospina | 0.03481866 | 200 km E of Rainbow | 0.2 µm | 2250 | STAPHYLOBACTIN |
| Bacteria ; Proteobacteria ; Gammaproteobacteria ; Enterobacterales ; Pseudoalteromonadaceae ; Pseudoalteromonas | 0.00939155 | 200 km E of Rainbow | 0.2 µm | 2250 | DESFERRIOXAMINE E , PUTREBACTIN / AVAROFERRIN , DESFERRIOXAMIN B , DESFERRIOXAMINE , UNKNOWN , STAPHYLOBACTIN |
| Bacteria ; Proteobacteria ; Gammaproteobacteria ; Pseudomonadales ; Pseudomonadaceae ; Pseudomonas | 0.0014487 | 200 km E of Rainbow | 0.2 µm | 2250 | XANTHOFERRIN , UNKNOWN , SYRINGAFACTIN , DESFERRIOXAMINE E , ARTHROFACTIN A , PUTREBACTIN / AVAROFERRIN , VIBRIOFERRIN , CICHOFACTIN A / CICHOFACTIN B , PUTISOLVIN , SYRINGOMYCIN , NUNAPEPTIN / NUNAMYCIN , COELICHELIN , DESFERRIOXAMIN B / DESFERRIOXAMINE E , CORPEPTIN A / CORPEPTIN B , ORFAMIDE A / ORFAMIDE C , ACINETOFERRIN , DESFERRIOXAMIN B , CAROTENOID , PYOVERDIN , GACAMIDE A , BANANAMIDE 1 / BANANAMIDE 2 / BANANAMIDE 3 |
| Bacteria ; Proteobacteria ; Gammaproteobacteria ; Enterobacterales ; Colwelliaceae ; Colwellia | 0.00084924 | 200 km E of Rainbow | 0.2 µm | 2250 | UNKNOWN |
| Bacteria ; Proteobacteria ; Gammaproteobacteria ; Pseudomonadales ; Marinobacteraceae ; Marinobacter | 0.00079928 | 200 km E of Rainbow | 0.2 µm | 2250 | PUTREBACTIN / AVAROFERRIN , XANTHOFERRIN , UNKNOWN |
| Bacteria ; Proteobacteria ; Gammaproteobacteria ; Enterobacterales ; Alteromonadaceae ; Alteromonas | 0.00014987 | 200 km E of Rainbow | 0.2 µm | 2250 | UNKNOWN |
| Bacteria ; Nitrospinota ; Nitrospinia ; Nitrospinales ; Nitrospinaceae ; Nitrospina | 0.09466131 | 200 km E of Rainbow | 0.2 µm | 600 | STAPHYLOBACTIN |

| | | | | | |
|---|---|---|---|---|---|
| Bacteria ; Proteobacteria ; Gammaproteobacteria ; Enterobacterales ; Pseudoalteromonadaceae ; Pseudoalteromonas | 0.0013824 | 200 km E of Rainbow | 0.2 µm | 600 | DESFERRIOXAMINE E , PUTREBACTIN / AVAROFERRIN , DESFERRIOXAMIN B , DESFERRIOXAMINE , UNKNOWN , STAPHYLOBACTIN |
| Bacteria ; Proteobacteria ; Gammaproteobacteria ; Enterobacterales ; Colwelliaceae ; Colwellia | 0.00125074 | 200 km E of Rainbow | 0.2 µm | 600 | UNKNOWN |
| Bacteria ; Nitrospinota ; Nitrospinia ; Nitrospinales ; Nitrospinaceae ; Nitrospina | 0.09409973 | 200 km E of Rainbow | 0.2 µm | 1600 | STAPHYLOBACTIN |
| Bacteria ; Nitrospinota ; Nitrospinia ; Nitrospinales ; Nitrospinaceae ; Nitrospina | 0.00867288 | Lucky Strike | 0.2 µm | 1670 | STAPHYLOBACTIN |
| Bacteria ; Proteobacteria ; Gammaproteobacteria ; Enterobacterales ; Pseudoalteromonadaceae ; Pseudoalteromonas | 0.00686845 | Lucky Strike | 0.2 µm | 1670 | DESFERRIOXAMINE E , PUTREBACTIN / AVAROFERRIN , DESFERRIOXAMIN B , DESFERRIOXAMINE , UNKNOWN , STAPHYLOBACTIN |
| Bacteria ; Proteobacteria ; Gammaproteobacteria ; Pseudomonadales ; Halomonadaceae ; Halomonas | 0.00314319 | Lucky Strike | 0.2 µm | 1670 | STAPHYLOBACTIN , UNKNOWN , DESFERRIOXAMINE E , PUTREBACTIN / AVAROFERRIN , AEROBACTIN , XANTHOFERRIN , VIBRIOFERRIN , BISUCABERIN B |
| Bacteria ; Proteobacteria ; Alphaproteobacteria ; Sphingomonadales ; Sphingomonadaceae ; Sphingobium | 0.00267753 | Lucky Strike | 0.2 µm | 1670 | UNKNOWN , XANTHOFERRIN , STAPHYLOBACTIN |
| Bacteria ; Proteobacteria ; Gammaproteobacteria ; Enterobacterales ; Vibrionaceae ; Vibrio | 0.0023865 | Lucky Strike | 0.2 µm | 1670 | VIBRIOFERRIN , PUTREBACTIN / AVAROFERRIN , AEROBACTIN , AMPHI-ENTEROBACTIN 1 / AMPHI-ENTEROBACTIN 2 / AMPHI-ENTEROBACTIN 3 / AMPHI-ENTEROBACTIN 4 , UNKNOWN |

| | | | | | |
|---|---|---|---|---|---|
| Bacteria ; Proteobacteria ; Gammaproteobacteria ; Pseudomonadales ; Marinobacteraceae ; Marinobacter | 0.00186263 | Lucky Strike | 0.2 µm | 1670 | PUTREBACTIN / AVAROFERRIN , XANTHOFERRIN , UNKNOWN |
| Bacteria ; Proteobacteria ; Alphaproteobacteria ; Rhodospirillales ; Thalassospiraceae ; Thalassospira | 0.00110594 | Lucky Strike | 0.2 µm | 1670 | UNKNOWN , STAPHYLOBACTIN |
| Bacteria ; Proteobacteria ; Gammaproteobacteria ; Enterobacterales ; Shewanellaceae ; Shewanella | 0.00069849 | Lucky Strike | 0.2 µm | 1670 | DESFERRIOXAMINE E , PUTREBACTIN / AVAROFERRIN , XANTHOFERRIN , UNKNOWN , VIBRIOFERRIN , STAPHYLOBACTIN |
| Bacteria ; Bacteroidota ; Bacteroidia ; Flavobacteriales ; Flavobacteriaceae ; Tenacibaculum | 0.00029104 | Lucky Strike | 0.2 µm | 1670 | BISUCABERIN B , UNKNOWN |
| Bacteria ; Proteobacteria ; Gammaproteobacteria ; Enterobacterales ; Pseudoalteromonadaceae ; Pseudoalteromonas | 0.02432581 | Lucky Strike | 3 µm | 1670 | DESFERRIOXAMINE E , PUTREBACTIN / AVAROFERRIN , DESFERRIOXAMIN B , DESFERRIOXAMINE , UNKNOWN , STAPHYLOBACTIN |
| Bacteria ; Proteobacteria ; Gammaproteobacteria ; Enterobacterales ; Shewanellaceae ; Shewanella | 0.01734722 | Lucky Strike | 3 µm | 1670 | DESFERRIOXAMINE E , PUTREBACTIN / AVAROFERRIN , XANTHOFERRIN , UNKNOWN , VIBRIOFERRIN , STAPHYLOBACTIN |
| Bacteria ; Proteobacteria ; Gammaproteobacteria ; Enterobacterales ; Moritellaceae ; Moritella | 0.01481233 | Lucky Strike | 3 µm | 1670 | COLICIN V , UNKNOWN |
| Bacteria ; Proteobacteria ; Gammaproteobacteria ; Enterobacterales ; Alteromonadaceae ; Alteromonas | 0.01284244 | Lucky Strike | 3 µm | 1670 | UNKNOWN |
| Bacteria ; Proteobacteria ; Alphaproteobacteria ; Sphingomonadales ; | 0.01071985 | Lucky Strike | 3 µm | 1670 | UNKNOWN , XANTHOFERRIN , STAPHYLOBACTIN |

| | | | | | |
|---|---|---|---|---|---|
| Sphingomonadaceae ; Sphingobium | | | | | |
| Bacteria ; Proteobacteria ; Gammaproteobacteria ; Enterobacterales ; Vibrionaceae ; Vibrio | 0.0050087 | Lucky Strike | 3 µm | 1670 | VIBRIOFERRIN , PUTREBACTIN / AVAROFERRIN , AEROBACTIN , AMPHI-ENTEROBACTIN 1 / AMPHI-ENTEROBACTIN 2 / AMPHI-ENTEROBACTIN 3 / AMPHI-ENTEROBACTIN 4 , UNKNOWN |
| Bacteria ; Proteobacteria ; Gammaproteobacteria ; Pseudomonadales ; Marinobacteraceae ; Marinobacter | 0.00464221 | Lucky Strike | 3 µm | 1670 | PUTREBACTIN / AVAROFERRIN , XANTHOFERRIN , UNKNOWN |
| Bacteria ; Bacteroidota ; Bacteroidia ; Flavobacteriales ; Flavobacteriaceae ; Flavobacterium | 0.00424518 | Lucky Strike | 3 µm | 1670 | BISUCABERIN B , PUTREBACTIN / AVAROFERRIN , UNKNOWN , DESFERRIOXAMINE E |
| Bacteria ; Proteobacteria ; Gammaproteobacteria ; Enterobacterales ; Colwelliaceae ; Colwellia | 0.00406194 | Lucky Strike | 3 µm | 1670 | UNKNOWN |
| Bacteria ; Nitrospinota ; Nitrospinia ; Nitrospinales ; Nitrospinaceae ; Nitrospina | 0.00404667 | Lucky Strike | 3 µm | 1670 | STAPHYLOBACTIN |
| Bacteria ; Proteobacteria ; Gammaproteobacteria ; Pseudomonadales ; Halomonadaceae ; Halomonas | 0.0040314 | Lucky Strike | 3 µm | 1670 | STAPHYLOBACTIN , UNKNOWN , DESFERRIOXAMINE E , PUTREBACTIN / AVAROFERRIN , AEROBACTIN , XANTHOFERRIN , VIBRIOFERRIN , BISUCABERIN B |
| Bacteria ; Proteobacteria ; Alphaproteobacteria ; Rhodobacterales ; Rhodobacteraceae ; Tateyamaria | 0.00299301 | Lucky Strike | 3 µm | 1670 | UNKNOWN |
| Bacteria ; Proteobacteria ; Gammaproteobacteria ; Coxiellales ; Coxiellaceae ; Coxiella | 0.00239746 | Lucky Strike | 3 µm | 1670 | UNKNOWN |

| | | | | | |
|---|---|---|---|---|---|
| Bacteria ; Proteobacteria ; Alphaproteobacteria ; Rhodobacterales ; Rhodobacteraceae ; Paracoccus | 0.00169502 | Lucky Strike | 3 µm | 1670 | UNKNOWN , CHEJUENOLIDE A / CHEJUENOLIDE B |
| Bacteria ; Actinobacteriota ; Actinobacteria ; Corynebacteriales ; Nocardiaceae ; Rhodococcus | 0.00169502 | Lucky Strike | 3 µm | 1670 | UNKNOWN |
| Bacteria ; Proteobacteria ; Gammaproteobacteria ; Pseudomonadales ; Spongiibacteraceae ; Sinobacterium | 0.00146596 | Lucky Strike | 3 µm | 1670 | PUTREBACTIN / AVAROFERRIN |
| Bacteria ; Proteobacteria ; Alphaproteobacteria ; Rhodospirillales ; Thalassospiraceae ; Thalassospira | 0.00071771 | Lucky Strike | 3 µm | 1670 | UNKNOWN , STAPHYLOBACTIN |
| Bacteria ; Proteobacteria ; Gammaproteobacteria ; Enterobacterales ; Vibrionaceae ; Photobacterium | 0.00062609 | Lucky Strike | 3 µm | 1670 | UNKNOWN , DESFERRIOXAMINE E , PUTREBACTIN / AVAROFERRIN , AEROBACTIN |
| Bacteria ; Proteobacteria ; Gammaproteobacteria ; Enterobacterales ; Psychromonadaceae ; Psychromonas | 0.00044284 | Lucky Strike | 3 µm | 1670 | UNKNOWN |
| Bacteria ; Proteobacteria ; Gammaproteobacteria ; Pseudomonadales ; Halomonadaceae ; Salinicola | 0.00042757 | Lucky Strike | 3 µm | 1670 | PUTREBACTIN / AVAROFERRIN , AEROBACTIN , DESFERRIOXAMINE E |
| Bacteria ; Spirochaetota ; Spirochaetia ; Spirochaetales ; Spirochaetaceae ; Treponema | 0.00029014 | Lucky Strike | 3 µm | 1670 | UNKNOWN |
| Bacteria ; Actinobacteriota ; Actinobacteria ; Propionibacteriales ; Nocardioidaceae ; Nocardioides | 0.00018325 | Lucky Strike | 3 µm | 1670 | DESFERRIOXAMIN B , DESFERRIOXAMINE , DESFERRIOXAMINE E |

| | | | | | |
|---|---|---|---|---|---|
| Bacteria ; Nitrospinota ; Nitrospinia ; Nitrospinales ; Nitrospinaceae ; Nitrospina | 0.02432 | 10 km S of Rainbow | 0.2 µm | 2000 | STAPHYLOBACTIN |
| Bacteria ; Proteobacteria ; Gammaproteobacteria ; Enterobacterales ; Pseudoalteromonadaceae ; Pseudoalteromonas | 0.00597333 | 10 km S of Rainbow | 0.2 µm | 2000 | DESFERRIOXAMINE E , PUTREBACTIN / AVAROFERRIN , DESFERRIOXAMIN B , DESFERRIOXAMINE , UNKNOWN , STAPHYLOBACTIN |
| Bacteria ; Proteobacteria ; Alphaproteobacteria ; Sphingomonadales ; Sphingomonadaceae ; Sphingobium | 0.00426667 | 10 km S of Rainbow | 0.2 µm | 2000 | UNKNOWN , XANTHOFERRIN , STAPHYLOBACTIN |
| Bacteria ; Proteobacteria ; Gammaproteobacteria ; Pseudomonadales ; Halomonadaceae ; Halomonas | 0.00202667 | 10 km S of Rainbow | 0.2 µm | 2000 | STAPHYLOBACTIN , UNKNOWN , DESFERRIOXAMINE E , PUTREBACTIN / AVAROFERRIN , AEROBACTIN , XANTHOFERRIN , VIBRIOFERRIN , BISUCABERIN B |
| Bacteria ; Proteobacteria ; Gammaproteobacteria ; Enterobacterales ; Alteromonadaceae ; Alteromonas | 0.00149333 | 10 km S of Rainbow | 0.2 µm | 2000 | UNKNOWN |
| Bacteria ; Bacteroidota ; Bacteroidia ; Flavobacteriales ; Flavobacteriaceae ; Flavobacterium | 0.00064 | 10 km S of Rainbow | 0.2 µm | 2000 | BISUCABERIN B , PUTREBACTIN / AVAROFERRIN , UNKNOWN , DESFERRIOXAMINE E |
| Bacteria ; Proteobacteria ; Gammaproteobacteria ; Enterobacterales ; Colwelliaceae ; Colwellia | 0.00032 | 10 km S of Rainbow | 0.2 µm | 2000 | UNKNOWN |
| Bacteria ; Proteobacteria ; Gammaproteobacteria ; Enterobacterales ; Alteromonadaceae ; Alteromonas | 0.11545231 | 10 km S of Rainbow | 3 µm | 2000 | UNKNOWN |
| Bacteria ; Proteobacteria ; Gammaproteobacteria ; Enterobacterales ; | 0.01854541 | 10 km S of Rainbow | 3 µm | 2000 | DESFERRIOXAMINE E , PUTREBACTIN / AVAROFERRIN , DESFERRIOXAMIN B , |

| | | | | | |
|---|---|---|---|---|---|
| Pseudoalteromonadaceae ; Pseudoalteromonas | | | | | DESFERRIOXAMINE , UNKNOWN , STAPHYLOBACTIN |
| Bacteria ; Proteobacteria ; Alphaproteobacteria ; Sphingomonadales ; Sphingomonadaceae ; Sphingobium | 0.01465153 | 10 km S of Rainbow | 3 µm | 2000 | UNKNOWN , XANTHOFERRIN , STAPHYLOBACTIN |
| Bacteria ; Proteobacteria ; Gammaproteobacteria ; Pseudomonadales ; Halomonadaceae ; Halomonas | 0.01333157 | 10 km S of Rainbow | 3 µm | 2000 | STAPHYLOBACTIN , UNKNOWN , DESFERRIOXAMINE E , PUTREBACTIN / AVAROFERRIN , AEROBACTIN , XANTHOFERRIN , VIBRIOFERRIN , BISUCABERIN B |
| Bacteria ; Proteobacteria ; Gammaproteobacteria ; Enterobacterales ; Moritellaceae ; Moritella | 0.00851373 | 10 km S of Rainbow | 3 µm | 2000 | COLICIN V , UNKNOWN |
| Bacteria ; Proteobacteria ; Gammaproteobacteria ; Enterobacterales ; Vibrionaceae ; Vibrio | 0.00714977 | 10 km S of Rainbow | 3 µm | 2000 | VIBRIOFERRIN , PUTREBACTIN / AVAROFERRIN , AEROBACTIN , AMPHI-ENTEROBACTIN 1 / AMPHI-ENTEROBACTIN 2 / AMPHI-ENTEROBACTIN 3 / AMPHI-ENTEROBACTIN 4 , UNKNOWN |
| Bacteria ; Nitrospinota ; Nitrospinia ; Nitrospinales ; Nitrospinaceae ; Nitrospina | 0.00688578 | 10 km S of Rainbow | 3 µm | 2000 | STAPHYLOBACTIN |
| Bacteria ; Proteobacteria ; Gammaproteobacteria ; Pseudomonadales ; Pseudomonadaceae ; Pseudomonas | 0.00400387 | 10 km S of Rainbow | 3 µm | 2000 | XANTHOFERRIN , UNKNOWN , SYRINGAFACTIN , DESFERRIOXAMINE E , ARTHROFACTIN A , PUTREBACTIN / AVAROFERRIN , VIBRIOFERRIN , CICHOFACTIN A / CICHOFACTIN B , PUTISOLVIN , SYRINGOMYCIN , NUNAPEPTIN / NUNAMYCIN , COELICHELIN , DESFERRIOXAMIN B / DESFERRIOXAMINE E , CORPEPTIN A / CORPEPTIN B , ORFAMIDE A / ORFAMIDE C , ACINETOFERRIN , DESFERRIOXAMIN B , CAROTENOID , PYOVERDIN , |

| | | | | | |
|---|---|---|---|---|---|
| | | | | | GACAMIDE A , BANANAMIDE 1 / BANANAMIDE 2 / BANANAMIDE 3 |
| Bacteria ; Bacteroidota ; Bacteroidia ; Flavobacteriales ; Flavobacteriaceae ; Flavobacterium | 0.00351989 | 10 km S of Rainbow | 3 µm | 2000 | BISUCABERIN B , PUTREBACTIN / AVAROFERRIN , UNKNOWN , DESFERRIOXAMINE E |
| Bacteria ; Proteobacteria ; Gammaproteobacteria ; Pseudomonadales ; Marinobacteraceae ; Marinobacter | 0.0031459 | 10 km S of Rainbow | 3 µm | 2000 | PUTREBACTIN / AVAROFERRIN , XANTHOFERRIN , UNKNOWN |
| Bacteria ; Proteobacteria ; Gammaproteobacteria ; Enterobacterales ; Vibrionaceae ; Photobacterium | 0.00277191 | 10 km S of Rainbow | 3 µm | 2000 | UNKNOWN , DESFERRIOXAMINE E , PUTREBACTIN / AVAROFERRIN , AEROBACTIN |
| Bacteria ; Proteobacteria ; Gammaproteobacteria ; Coxiellales ; Coxiellaceae ; Coxiella | 0.00217793 | 10 km S of Rainbow | 3 µm | 2000 | UNKNOWN |
| Bacteria ; Proteobacteria ; Gammaproteobacteria ; Enterobacterales ; Colwelliaceae ; Colwellia | 0.00178194 | 10 km S of Rainbow | 3 µm | 2000 | UNKNOWN |
| Bacteria ; Proteobacteria ; Alphaproteobacteria ; Sphingomonadales ; Sphingomonadaceae ; Sphingomonas | 0.00167195 | 10 km S of Rainbow | 3 µm | 2000 | STAPHYLOBACTIN |
| Bacteria ; Proteobacteria ; Gammaproteobacteria ; Pseudomonadales ; Halomonadaceae ; Salinicola | 0.00145195 | 10 km S of Rainbow | 3 µm | 2000 | PUTREBACTIN / AVAROFERRIN , AEROBACTIN , DESFERRIOXAMINE E |
| Bacteria ; Proteobacteria ; Gammaproteobacteria ; Enterobacterales ; Shewanellaceae ; Shewanella | 0.00118796 | 10 km S of Rainbow | 3 µm | 2000 | DESFERRIOXAMINE E , PUTREBACTIN / AVAROFERRIN , XANTHOFERRIN , UNKNOWN , VIBRIOFERRIN , STAPHYLOBACTIN |
| Bacteria ; Proteobacteria ; Gammaproteobacteria ; | 0.00107797 | 10 km S of Rainbow | 3 µm | 2000 | DESFERRIOXAMINE E |

| | | | | | |
|---|---|---|---|---|---|
| Pseudomonadales ; Saccharospirillaceae ; Oleibacter | | | | | |
| Bacteria ; Proteobacteria ; Gammaproteobacteria ; Francisellales ; Francisellaceae ; Fangia | 0.00107797 | 10 km S of Rainbow | 3 µm | 2000 | UNKNOWN |
| Bacteria ; Proteobacteria ; Gammaproteobacteria ; Pseudomonadales ; Spongiibacteraceae ; Sinobacterium | 0.00061598 | 10 km S of Rainbow | 3 µm | 2000 | PUTREBACTIN / AVAROFERRIN |
| Bacteria ; Proteobacteria ; Gammaproteobacteria ; Enterobacterales ; Psychromonadaceae ; Psychromonas | 0.00048398 | 10 km S of Rainbow | 3 µm | 2000 | UNKNOWN |
| Bacteria ; Proteobacteria ; Alphaproteobacteria ; Rhodobacterales ; Rhodobacteraceae ; Paracoccus | 0.00046199 | 10 km S of Rainbow | 3 µm | 2000 | UNKNOWN , CHEJUENOLIDE A / CHEJUENOLIDE B |
| Bacteria ; Proteobacteria ; Gammaproteobacteria ; Burkholderiales ; Oxalobacteraceae ; Massilia | 0.00035199 | 10 km S of Rainbow | 3 µm | 2000 | CUPRIACHELIN , UNKNOWN , XANTHOFERRIN |
| Bacteria ; Proteobacteria ; Gammaproteobacteria ; Pseudomonadales ; Moraxellaceae ; Psychrobacter | 0.00024199 | 10 km S of Rainbow | 3 µm | 2000 | VIBRIOFERRIN , ACINETOFERRIN |

Solid lines separate the three different stations. Dashed lines separate 0.2 µm and 3 µm filter sizes investigated in this study.

St. 7 = Lucky Strike
St. 11 = 200 km E of Rainbow
St. 17 = 10 km S of Rainbow

See **Supplementary Table 1** for additional information.
